# Disentangling the biotic and abiotic drivers of bird-building collisions in a tropical Asian city using ecological niche modeling

**DOI:** 10.1101/2023.06.27.546782

**Authors:** David J. X. Tan, Nicholas A. Freymueller, Kah Ming Teo, William S. Symes, Shawn K.Y. Lum, Frank E. Rheindt

## Abstract

Bird-building collisions are responsible for a large number of bird deaths in cities around the world, yet they remain poorly studied outside of North America. This study presents one of the first city-wide fine-scale and landscape-scale analyses of bird-building collisions from Asia and represents a novel application of maximum entropy modeling (as commonly applied to species distribution modeling) to assess the drivers of bird-building collisions in the tropical city-state of Singapore. Our results show that the drivers of bird-building collisions often vary among taxa, with several migratory taxa having a higher relative collision risk linked to areas with high building densities and high levels of nocturnal blue light pollution. In contrast, non-migratory taxa had a higher collision risk in areas proximate to woodland cover. Projecting these models onto high-fidelity long-term government land-use plans, we demonstrate that our approach can be applied to predict future changes in bird-building collision risk stemming from future increases in blue light pollution and encroachment of buildings into forested areas. Our results suggest that bird-building collision mitigation measures need to account for the differential drivers of collision across both resident and migratory species, and show that combining community science and ecological modeling can be a powerful approach for analyzing bird-building collision data.

**Article impact statement:** Inferring the drivers and distribution patterns of bird-building collision hotspots in Singapore using community science and maximum entropy modeling

## Introduction

Bird-building collisions are prevalent across cities worldwide and represent a significant source of urban avian mortalities (Klem, 1989; Loss et al.,, 2015). In North America alone, it is estimated that between 365 to 988 million birds die from building collisions annually (Loss et al., 2014), second only to the number of cat predation-related mortalities (Loss et al., 2013).

Despite the global nature of this phenomenon, significant geographical and methodological biases exist in our understanding of bird-building collisions. For one, most bird-building collision studies have tended to focus on temperate North America, specifically on migratory Nearctic species (Basilio et al., 2020), with relatively few studies focused on the tropics, especially the Afro- and Oriental tropics (Basilio et al., 2020; Tan et al., 2017). Furthermore, most studies of bird-building collisions have thus far been based on systematic surveys conducted at very fine spatial scales, with survey transects ranging from 1 to 21 buildings, usually on university campuses or within city centers (Hager et al., 2017; Elmore et al., 2020). While these studies broadly suggest that collision rates are correlated with urban abiotic factors such as the glass area on building façades (Hager et al., 2008; Borden et al., 2010; Cusa et al., 2015), building size (Hager et al., 2017; Elmore et al., 2020), nocturnal light pollution (Longcore & Rich, 2004; Winger et al., 2019; Lao et al., 2020), and regional urbanization (Hager et al., 2017), as well as biotic factors such as vegetation density (Cusa et al., 2015) and vegetation proximity (Gelb & Delacretaz, 2009; Barton et al., 2017; Rebolo-Ifrán et al., 2019), the limited scale at which these surveys have been conducted means they are unlikely to capture the full heterogeneity of city landscapes (including both urban and peri-urban areas). Furthermore, extending such fine-scale surveys to a city-wide scale is often unfeasible. Alternative surveying strategies based on opportunistic community/citizen science approaches (Bayne et al., 2012; Rebolo-Ifrán et al., 2019) are often difficult to quantitatively analyze owing to the lack of true absence data. Consequently, our understanding of how bird-building collision risk varies across geographical space, at local, regional, and global scales, is limited.

The city-state of Singapore has a well-documented incidence of bird-building collisions (Low et al., 2017; Tan et al., 2017), likely owing to its high level of urbanization (United Nations, 2018) and landscape heterogeneity (Gaw et al., 2019). Even so, little is known about the driving factors determining the spatial pattern of bird-building collisions in Singapore. In addition, while community science observations have identified resident and migratory bird species that may be particularly susceptible to building collision mortalities (e.g. Pink-necked Green Pigeon (*Treron vernans*), Blue-winged Pitta (*Pitta moluccensis*)), it remains unclear if the factors driving collisions in Singapore differ among species (Elmore et al., 2020; Tan et al., 2017).

One useful analytical approach that has largely been overlooked in the context of bird-building collisions is maximum entropy modeling, specifically MaxEnt (Phillips et al., 2006), which quantifies the relative probability of occurrence events by comparing known presence-only occurrence records against a background of environmental predictor variables. Although MaxEnt is more often used for modeling the distributions of living organisms, it is conceptually equivalent to apply MaxEnt to modeling the distribution of bird-building mortalities, as the risk of bird-building collisions can be similarly described as a stochastic Poisson phenomenon with a probability density dependent on environmental covariates that are inferred to have a causal relationship with occurrence/mortality records (Elith et al., 2010). The ability of MaxEnt to model species distributions based on presence-only occurrences makes this method particularly suitable for analyzing unstructured community science data so long as sampling bias is accounted for. Maximum entropy modeling and other species distribution modeling-adjacent techniques have already been applied to understand causes of mortality in birds, including roadkill (Gomes et al., 2008), power lines, and wind turbines (Smeraldo et al., 2020), while leveraging citizen-science data.

Here, we address these gaps and methodological challenges in our understanding of bird-building collisions by analyzing a multi-year community science dataset of bird-building collisions from Singapore through a novel application of maximum entropy models. We use MaxEnt to parameterize relative collision risk models for both resident and migratory bird species and use these models to predict likely bird-building collision hotspots across the island of Singapore. We subsequently project these models onto future conditions based on high-fidelity land-use plans, and predict how collision risks in Singapore are likely to change over the next 10 to 15 years. In so doing, we demonstrate how MaxEnt-based analyses can be a powerful tool for modeling bird-building collision risks, which will be useful to both urban planners and conservationists alike.

## Methods

### Study Area and Design

This study was conducted in the island nation of Singapore (1.1°-1.4°N and 103.6°-104.1°E; ∼ 725 km^2^), which is located within the Sundaland biodiversity hotspot as well as along the East Asian-Australasian Flyway (Fig. 1). We documented bird-building collision records in Singapore from 2013 to 2020 using an established community science approach by encouraging members of the public to report sightings of dead birds to a public hotline as well as via social media (Low et al., 2017). We identified all reported sightings to species based on photographic documentation or, where possible, collected specimens, and georeferenced all bird mortality reports where possible based on GPS coordinates obtained from Google Maps. We assessed the likely cause of injury or death based on several lines of evidence as described in Tan et al. (2017). Specifically, birds found at the base of buildings with signs of facial injury or head trauma were classified as building collision victims. Other forms of mortality were also documented, and birds for which the cause of death could not be identified were listed as ‘unknown’ (Tan et al., 2017). Based on the bird-building records collected, we collated a database of confirmed bird-building collision records for which locality data was available, and classified each record based on the species’ migratory status, excluding introduced species and species for which the migratory status is indeterminate. Finally, we spatially thinned occurrences to a single record per taxon per grid cell.

**Figure 1:**
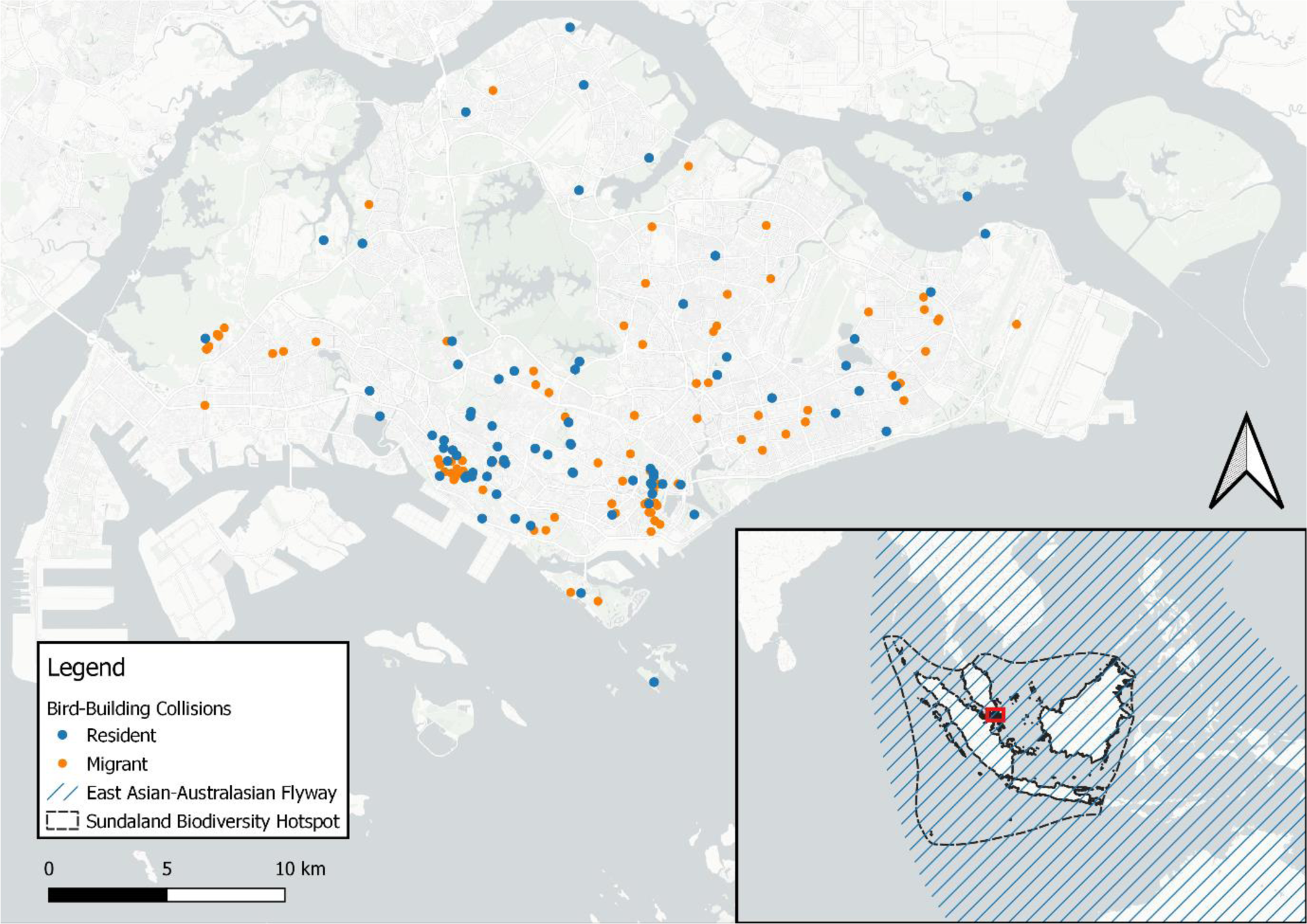
Bird-building collision localities in Singapore. Blue collisions involved non-migratory species and orange collisions involved migratory species. A total of 224 confirmed collision records were documented from 2013 to 2020. Inset depicts Singapore’s location relative to South and Southeast Asia, showing the island’s position within the Sundaland biodiversity hotspot and the East Asian-Australasian Flyway

We modeled responses of specific avian taxa by pooling phylogenetically similar species into taxonomic bins in order to increase sample sizes for each subset, and prepared subsets of the full collision dataset for taxa with at least 10 confirmed collision records. These taxon bins include migratory taxa such as pittas (genus *Pitta*), bitterns (genus *Ixobrychus*), *Ficedula* flycatchers (genus *Ficedula*), and the Black-backed Kingfisher (*Ceyx erithacus*), and non-migratory taxa such as green pigeon (genus *Treron*) and the Asian Emerald Dove (*Chalcophaps indica*). In addition, we also prepared a separate subset of collision records for all migrants and all residents (Table 1).

**Table 1:**
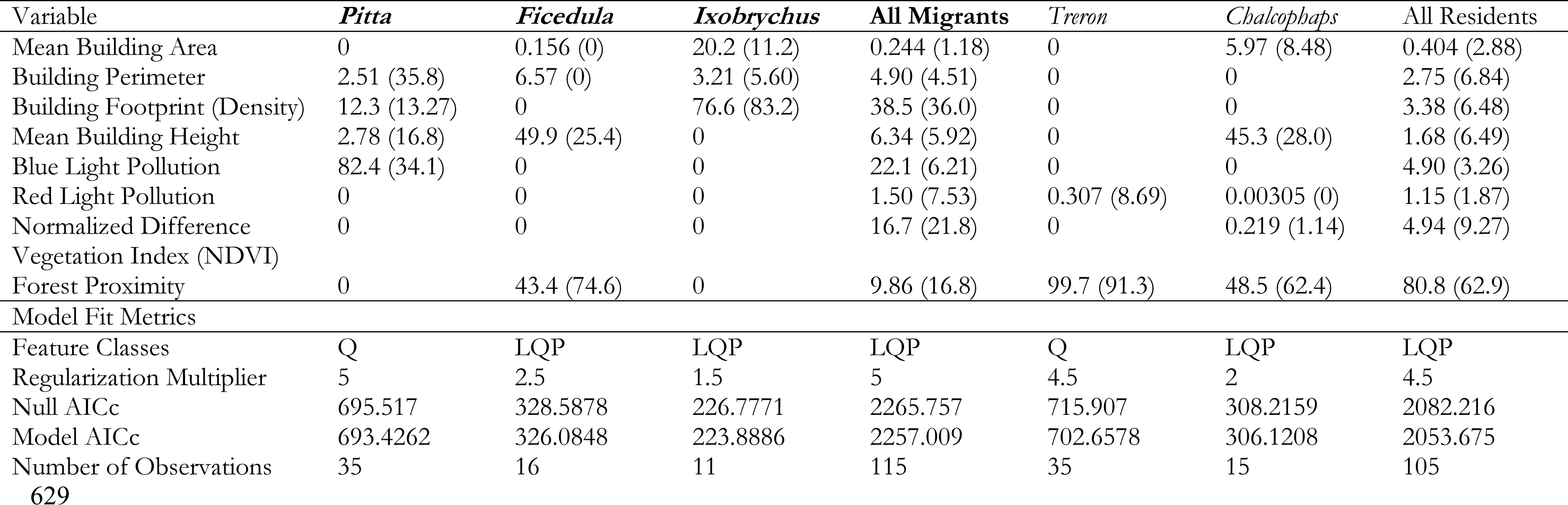
Best fit fine-scale (100 m x 100 m) model loadings for each taxon group/migratory phenology based on the Akaike’s Information Criterion corrected for small sample sizes (AICc), with variable percentage contribution and permutation importance (in parentheses) in the upper table, and model fit metrics reported in the lower table. Taxon names in bold are migratory, while non-bolded taxa are non-migratory.

### Abiotic Environmental Predictors

We built raster layers based on the spatial urban environment by downloading building polygons from the OpenStreetMap database (number of buildings = 104,393) to model the effect of building size and density on bird-building collisions. OpenStreetMap contains a near-complete set of building polygons for Singapore (Biljecki, 2020) in shapefile format. We combined buildings with shared edges via the ‘Dissolve’ and ‘Multipart to Singleparts’ functions in QGIS v3.14.16 since birds are unlikely to perceive subdivided buildings as separate entities. From the simplified building dataset (number of buildings = 72,448), we generated a 100 x 100 meter rectangular grid over the map extent and used the ‘Join Attributes by Location’ function in QGIS to extract building attribute summaries. To calculate the per pixel building perimeter, we used the ‘Polygons to Lines’ function in QGIS to convert the building polygon dataset into a series of lines and calculated the total line length contained within each 100 x 100m pixel. We generated rasters of building size and density by calculating the total within-pixel building footprint as a measure of building density, as well as the mean floor area of all buildings intersecting each pixel as a measure of building size. To generate a raster of building heights, we manually annotated building polygons from the full OpenStreetMap dataset with building height data from Biljecki (2020) and OneMap 3.0 (https://www.onemap3d.gov.sg/main/, Singapore Land Authority), excluding sensitive structures such as government buildings and military installations. We subsequently coarsened the dataset by applying a 100 x 100-meter rectangular grid over the map extent and used the “Join Attribute by Location” function in QGIS to calculate per-pixel mean building heights.

We quantified the effects of nighttime light pollution by downloading a true color night photograph of Singapore taken using a Nikon D4 with a 400 mm telephoto lens on 17 March 2016 by astronaut Tim Kopra from the International Space Station (NASA, 2016), and georeferenced the image in QGIS. We subsequently decomposed the image into its red, blue, and green color channels to isolate the effect of different light spectra on bird-building collisions. We resampled all nighttime light pollution rasters using the *r.resamp.interp()* function in GRASS to a 100 m resolution using a bilinear interpolation.

### Biotic Environmental Predictors

To generate rasters of vegetation density, we calculated the normalized difference vegetation index (NDVI) from a cloud-free multispectral image of Singapore captured by the LandSat8 OLI/TIRS platform (USGS). In addition, we applied the ‘Proximity (Raster Distance)’ function in QGIS to a rasterized map of Singapore’s woodland cover in 2013 (Tan et al., 2018) to generate a forest proximity raster. These rasters were subsequently resampled to 100 m resolution using the GRASS GIS *r.resamp.interp()* function using a bilinear interpolation method.

All predictor variable rasters and input shapefiles generated for this analysis were reprojected to the SVY21/Singapore TM projection (EPSG:3414). The complete list of predictor variables used in this study can be found in Table 1 and Figure 2.

**Figure 2:**
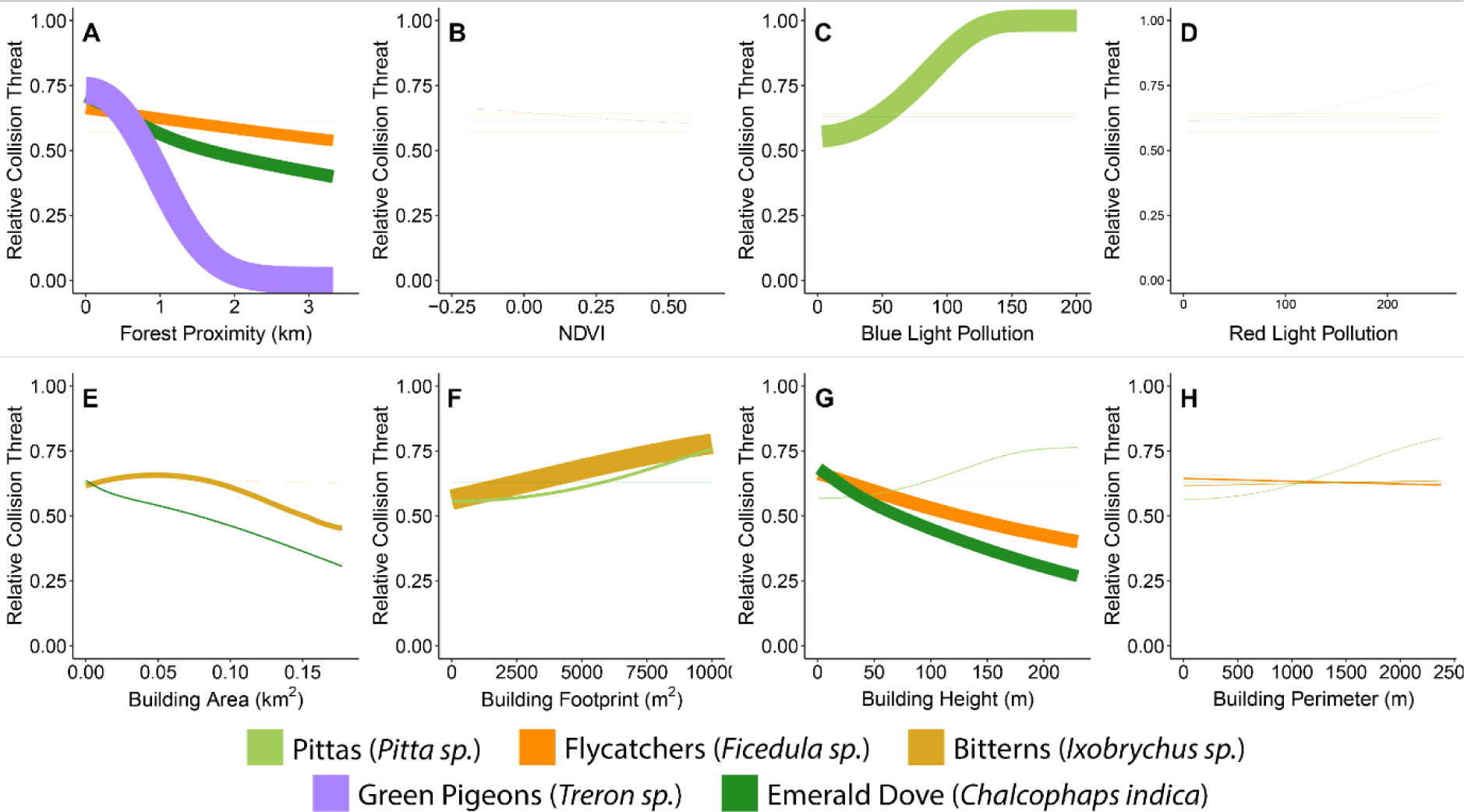
Taxon-specific response curves for all eight fine-scale ecogeographic predictor variables: (A) Forest proximity, (B) Normalized Difference Vegetation Index (NDVI), (C) nocturnal blue light pollution, (D) nocturnal red light pollution, (E) mean per-pixel building area, (F) total per-pixel building footprint (i.e. the area of a 100m x 100m pixel occupied by buildings; an estimator of building density), (G) mean per-pixel building height, and (H) total per-pixel building perimeter. The color of each line corresponds to the respective avian taxon, and the thickness of each line indicates the percentage contribution of the variable to the respective taxon-specific model.

### Calculating coarse-scale landscape effects

To investigate how collision risk varies with spatial scale, we generated coarsened versions of our predictor variable dataset using the GRASS GIS *r.neighbors()* function, which resamples pixel values within a 500 m circular neighborhood. We generated coarsened focal rasters for all predictor variables except forest proximity, which allowed us to test the sensitivity of bird-building collisions at different spatial scales.

### Creating future land-use change realization scenarios

To model how the distributions of bird-building collisions are expected to shift with land-use change, we generated a set of future land-use scenarios based on the 2019 Singapore Master Plan (Urban Redevelopment Authority, 2019), which maps out high-fidelity land-use strategies in Singapore for the next 10-15 years. We generated a simplified list of 11 land-use types based on the 2019 Master Plan (Appendix S1) and identified a subset of baseline polygons corresponding to areas where land-use change was not planned, as well as a separate subset of polygons corresponding to areas earmarked for land-use change. We used QGIS to extract baseline values for each predictor variable and land-use type combination, and used the function *fitdist()* in the R package *fitdistrplus* (Delignette-Muller & Dutang, 2015) to fit a gamma distribution for each combination of land-use type and predictor variable using the maximum likelihood estimator method (Appendix S1). We used the gamma distribution since the predictor variables generally exhibit positive skewed distributions. With parameters obtained from the fitted gamma distributions, we used the ‘Create Random Raster Layer (Gamma Distribution)’ function in QGIS to generate a random raster for each combination of predictor variable and land-use type (Appendix S1), clipped to the predicted land-use extent based on the 2019 Singapore Master Plan, for five independent replicates. For predictor variables with negative (e.g. NDVI) or very high pixel values (e.g. mean building area), we applied a transformation to the baseline raster values to facilitate gamma distribution fitting (Appendix S1), and subsequently corrected the transformation in the randomized predicted land-use rasters. We generated an overall land-use change grid for each independent replicate by using the *r.series()* function in GRASS GIS, which combines the predicted grids for each land-use category into a single output grid, and averages the grid value for overlapping grids with multiple land-use types. We used the raster calculator in QGIS to mask pixels in the original predictor variable rasters that intersect with the land-use change grids and overlaid the land-use change grids over the original predictor variable rasters.

Singapore is transitioning from high-pressure sodium vapor streetlamps (which emit red/orange light) to LED streetlights (which emit white light; Land Transport Authority, 2017). We accounted for this transition by extracting present-day streetlight pollution values from the red-light pollution raster using a shapefile of Singapore’s road network and adding varying proportions of present-day red-wavelength streetlight pixel values to the blue light pollution raster. This is based on the assumption that future LED streetlights will be maintained at a similar level of brightness as present-day sodium vapor streetlamps. We modelled five future blue light pollution scenarios ranging from 20% to 100% of present-day red light pollution levels (in increments of 20%) and generated five randomized future blue light pollution rasters per scenario based on the methods described above.

### Predictor Variable Thinning

To test for predictor variable collinearity, we used the *r.covar()* GRASS GIS function to calculate a covariance matrix for all possible pairwise comparisons of predictor variables, and excluded the total building area, green light pollution, coarse-scale green light pollution, and coarse-scale mean building area predictors due to their high correlations with other predictor variables (R > 0.8; Appendix S2). We clipped the predictor variable rasters to exclude areas where collision detection rates were likely to be anomalously low, including: military areas; nature reserves; high security industrial zones; and Changi Airport. This resulted in 26,821 background points.

### Model Generation

We generated maximum entropy (MaxEnt) models v3.4.1 (Phillips et al., 2017) using a pipeline based on the *dismo* v1.3-5 and kuenm v0.1.1 R packages (Hijmans et al., 2015; Cobos et al., 2019) as described in Freymueller (2020). We ran MaxEnt models with ‘Q’ and ‘LQP’ feature class combinations and regularization multiplier (β) values from 1 to 5. To assess model fit, we used the Akaike Information Criterion metric adjusted for small sample sizes (AICc; sensu Warren & Seifert, 2011) and selected models with the lowest AICc values (ΔAICc_minimum_ < 2) and a ΔAICc relative to a random null model greater than 2. We applied this non-traditional MaxEnt approach to ensure that our models were not unnecessarily complex/overfit, especially given that some species had low sample sizes. After identifying the most likely models for each species, we visually inspected the response curves to ensure that they displayed ecologically plausible (i.e., concave-down) behavior. Any models that displayed concave-up response curves were re-run after excluding problem variables from analysis. Whenever multiple models emerged with high support (ΔAICc_minimum_ < 2), we compared them using five-fold cross-validation and selected the model that had lower intra-model variability. We similarly performed such cross-validation on models that emerged as the sole “best-supported” model to ensure they were not sensitive to slight changes in input data, which would suggest that they are overfit and not informative for model projection to new environmental conditions (Peterson and Samy 2016). Following cross-validation, we generated weighted averages of all five folds into an ensemble model for each species. Weights were determined by multiplying testing sensitivity (given a training threshold of lowest presence; sensu Pearson et al., 2006) by the partial area under the receiver operating characteristic curve (partialROC; at a defined omission of 0.01 across 500 iterations; Peterson et al., 2008). Testing sensitivity was appropriate to include as part of the model weights as it exists independent of any absence/fractional predicted area metric. We calculated partialROC metrics using the kuenm R package (Cobos et al., 2019); partialROC is more appropriate than traditional ROC-AUC-based metrics for correlative MaxEnt modeling given the lack of true absence data (Peterson et al., 2008; Cobos et al., 2019).

We ran MaxEnt models for a total of six avian taxa (pittas (*Pitta* sp.), bitterns (*Ixobrychus* sp.), *Ficedula* flycatchers, Black-backed Kingfisher (*Ceyx erithaca*), green pigeons (*Treron* sp.), and Asian Emerald Dove (*Chalcophaps indica*)), as well as two additional models containing all migratory and all non-migratory birds (Table 1 and Appendix S3). For all taxa, we ran one set of models using the fine-scale predictor variables (100m resolution) and the other with the landscape-scale predictors (500m resolution). We excluded Asian Glossy Starlings (*Aplonis panayensis*) from this analysis due to the relatively small number of occurrence records (N=6) retained post spatial thinning. We subsequently projected the best models from the 100m resolution dataset onto the full unclipped set of predictor variable rasters and plotted collision risk maps using the *ggplot2* R package (Wickham, 2016; Fig. 3; Appendices S5A, S5C).

**Figure 3:**
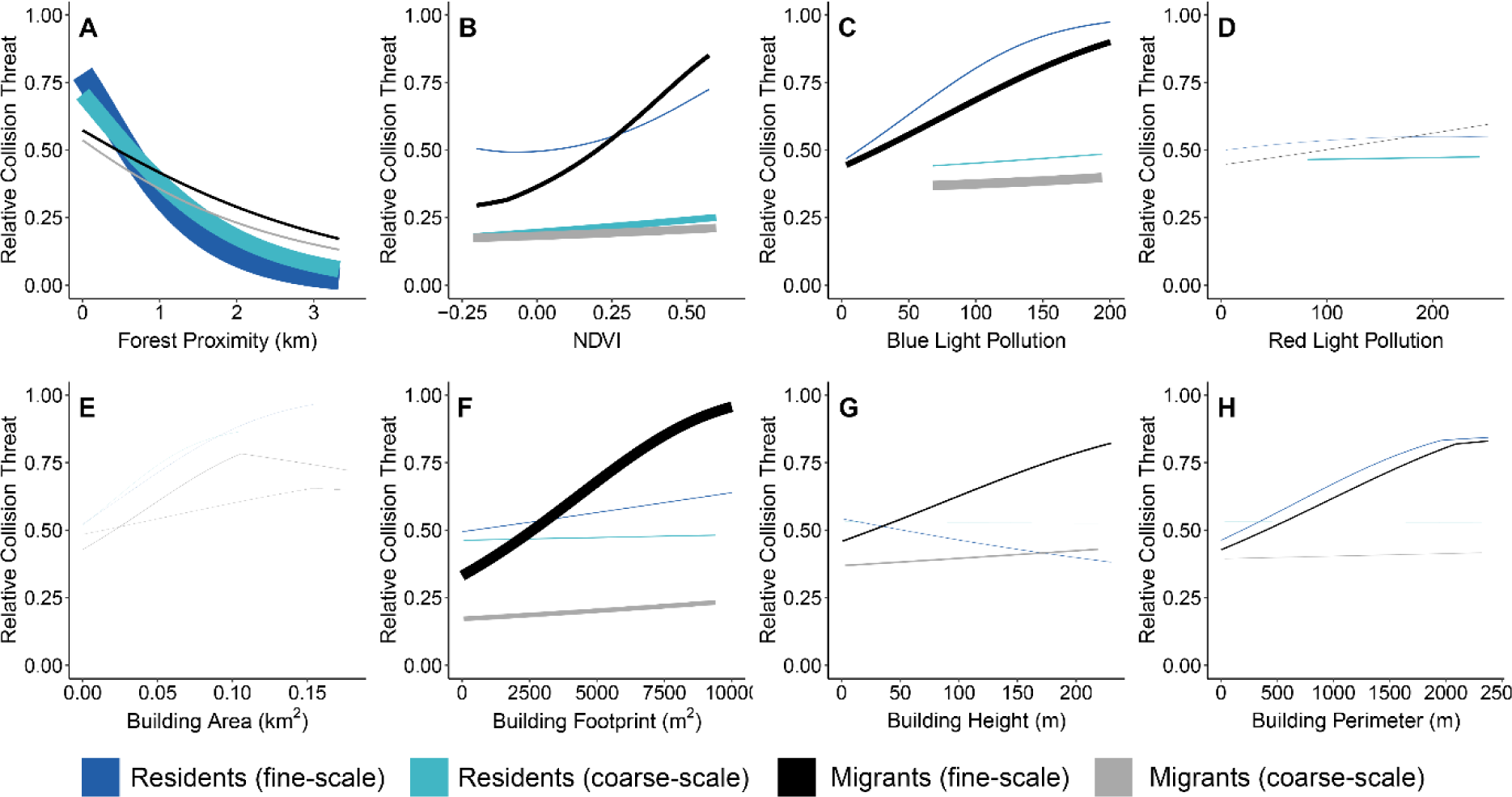
Response curve plots for different migratory phenologies and spatial scales, with line thickness corresponding to the percentage contribution of the variable to the respective model. The results indicate that, regardless of spatial scale, forest proximity is the strongest contributor to resident bird collisions in Singapore, while Normalized Difference Vegetation Index (NDVI), nocturnal blue light pollution, and building footprint (i.e. building density) are the strongest predictors of migratory bird collisions in Singapore.

We generated future collision risk maps for each taxon by projecting the best model for each taxon onto all five predictor variable replicates for each of the five blue light pollution scenarios, for a total of 25 future collision risk maps. We generated a final forecast map for each taxon and blue light pollution scenario by averaging over the five replicates and plotted the maps using ggplot2 (Figs. 5, 6; Appendices S6, S7). We ensured that these future projections were limited to areas with positive Multivariate Environmental Similarity Surface (MESS; Elith et al., 2010) values to avoid making predictions in non-analogous environmental space, where the transferability of our models may be diminished (Appendices S9-S16). All areas contained positive MESS values.

**Figure 4:**
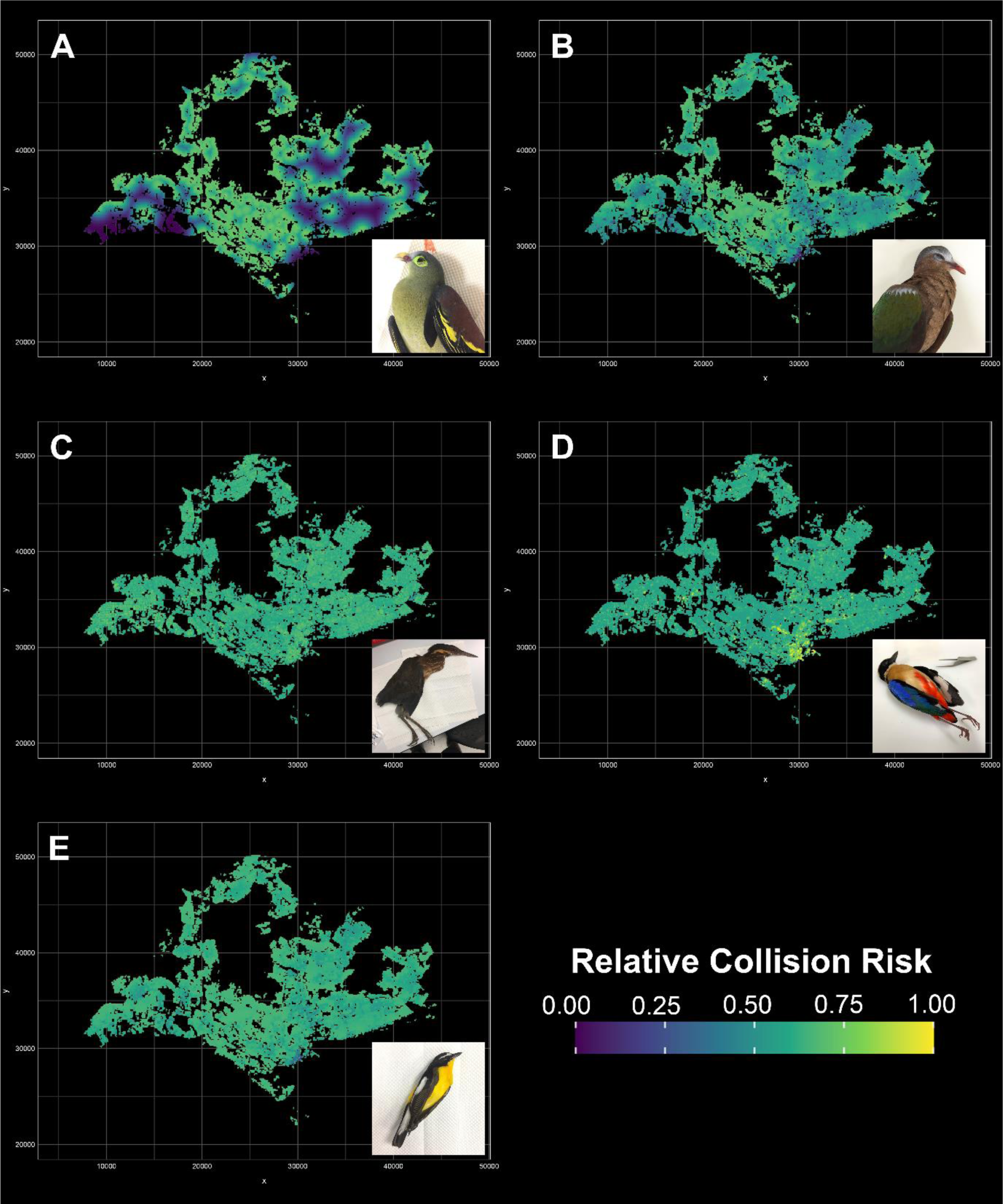
Building collision risk maps for five “supercollider” taxa in Singapore: (A) green pigeons (*Treron sp.*), (B) Asian Emerald Dove (*Chalcophaps indica*), (C) bitterns (*Ixobrychus sp.*), (D) pittas (*Pitta sp.*), and (E) *Ficedula* flycatchers (*Ficedula sp.*) based on fine-scale predictor variables, showing a high landscape-wide collision risk for all taxa, and a pronounced collision risk zone for pittas in the downtown area of Singapore. Note that these collision risk values are relative, and a collision risk of 1 does not equate to a 100% risk of collision.

**Figure 5:**
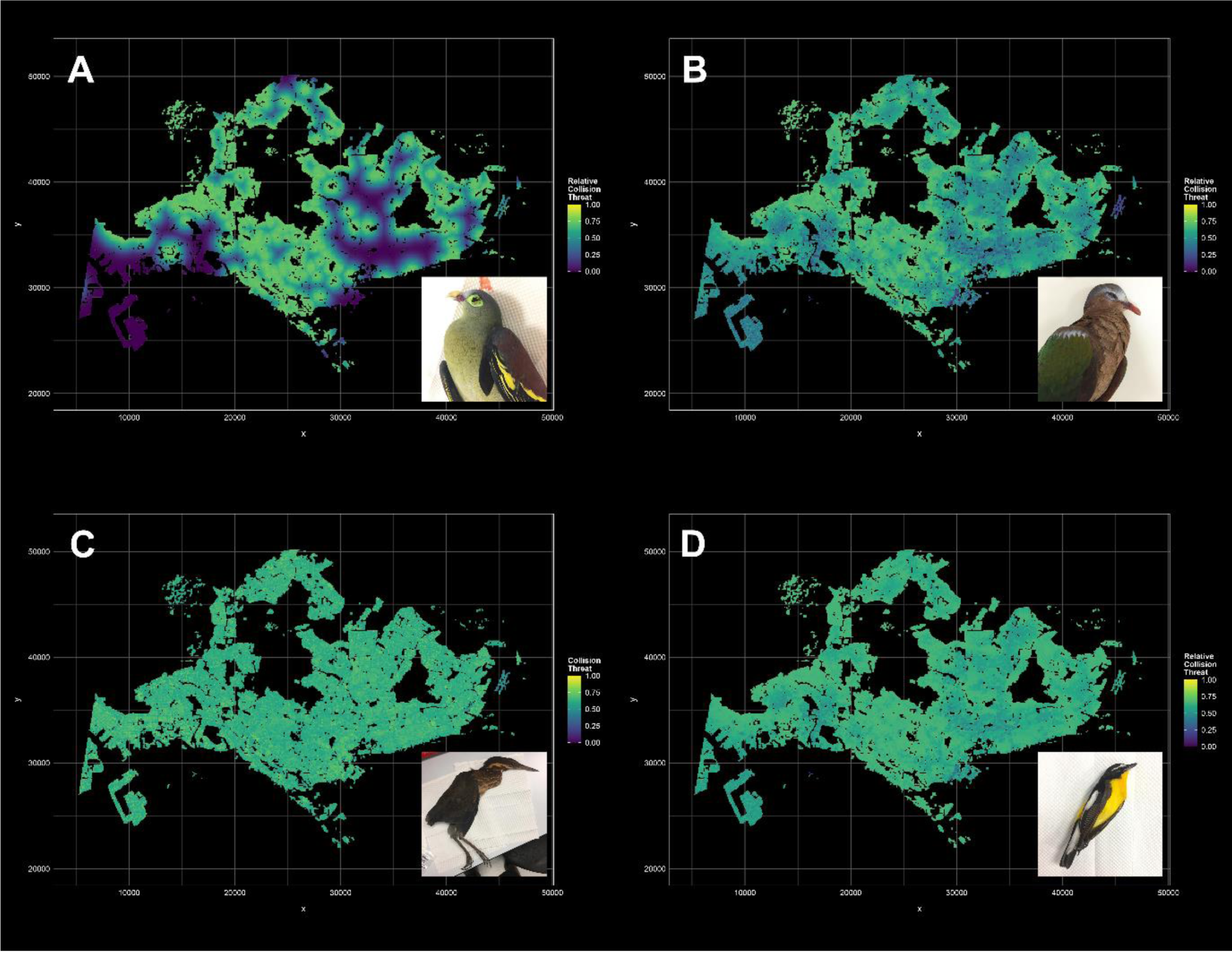
Projected future fine-scale collision risk maps for (A) green pigeons (*Treron sp.*), (B) Asian Emerald Dove (*Chalcophaps indica*), (C) bitterns (*Ixobrychus sp.*), and (D) *Ficedula* flycatchers (*Ficedula sp.*) based on the Singapore Master Plan 2019, showing that future urban developments are likely to pose a high collision risk to both migratory and resident birds, especially as the fragmentation of extant forest creates new forest edges.

## Results

### Bird-Building Collision Records

We compiled a total of 224 confirmed bird-building collision records between 2013 and 2020 (mean of 34.3 detected collisions per year, ranging from 12 to 51 observations annually), of which 115 were migrants and 105 were residents (Fig. 1). Of the migrants recorded, 63.4% of collisions were comprised of only 8 species (24% of total species richness), with 35 records (30.4%) from the genus *Pitta* (*P. moluccensis* (27) and *P. sordida* (8)), 16 records (13.9%) from the genus *Ficedula* (*F. zanthopygia* (15) and *F. mugimaki* (1)), 11 (9.57%) from the genus *Ixobrychus* (*I. flavicollis* (5), *I. cinnamomeus* (5), and *I. sinensis* (1)), and 11 (9.57%) of the Black-backed Kingfisher. As for resident species, collision mortalities were dominated by pigeons (47.6%), with 33 Pink-necked Green Pigeon (*Treron vernans*), 2 Thick-billed Green Pigeon (*Treron curvirostra*) and 15 Asian Emerald Dove records.

### MaxEnt Model Outputs

The variable coefficients and response curves of our MaxEnt models indicate that for each taxon group, collision risk can be explained by two to three high-contributing variables, and that these high-contributing variables differ among taxa (Fig. 2). In more general terms, our MaxEnt models suggest that forest proximity (Figs. 2A, 3A) may be one of the most important predictors of bird-building collisions in the Southeast Asian tropics, affecting both resident and some migratory taxa. In particular, our taxon-specific models suggest that low to medium-rise buildings located near forest edge appear to be collision hotspots for both emerald doves and *Ficedula* flycatchers (Figs. 4B and 4E). Although building size (as quantified by both building perimeter and mean building area; Figs. 2E, 2H, 3E, and 3H) does to some extent influence bird-building collision risk in bitterns and emerald doves, its effect is generally weaker relative to other predictor variables, contrary to prior studies on bird-building collisions (Hager et al., 2017; Elmore et al., 2020). Similarly, red light pollution (Figs. 2D and 3D) appears to have a considerably lesser effect on collision risk as compared to blue light pollution, which points to the differential effects of specific light spectra on avian behavior. We did not observe any strong species-specific effect of vegetation density on collision risk, and only observed a correlation between NDVI and collision risk in analyses broadly merging all migrants and, at landscape scales, non-migrants (Fig. 2B and 3B).

Across all migrants, the best-performing model indicated that building density was the strongest predictor of building collisions (38.5%), followed by blue light pollution (22.1%) and vegetation density (16.7%) at the fine scale (Table 1; Fig. 3). The same variables were also identified in the landscape-scale model, although blue light pollution was the strongest predictor (34.8%) followed by vegetation density (30.3%) and building density (17.6%) (Appendix S3; Fig. 3). For the pittas in particular, high blue light pollution was the strongest predictor of collision at both fine and landscape scales (82.4% and 78.2%, respectively; Tables 1 & S3). For bitterns, high building density and building size were the main predictors of collisions, with fine-scale building footprint and mean building area explaining 76.6% (Fig. 2F) and 20.2% (Fig. 2E) of the best model, respectively (Table 1), while at landscape scales the best model did not emerge as more likely than the null model (Appendix S3). Unlike for other migratory taxa, the best performing model for *Ficedula* flycatcher collision risk encompassed forest proximity and low building height (43% and 49.9% respectively; Table 1), with no clear pattern at coarser spatial scales (Appendix S3). We were unable to obtain any useful explanatory models for Black-backed Kingfishers, likely due to small sample size (n=11).

Across resident species, the best-performing model at the fine scale identified forest proximity as the strongest predictor of bird-building collisions (80.8%), with all other predictors having <10% contribution (Fig. 3). At the landscape scale, forest proximity equally emerged as the best predictor of collision risk (59.5%) across resident birds, although NDVI also appeared to be a strong predictor (25.6%) (Fig. 3). Both the best performing fine and landscape scale models for *Treron* green pigeons identified forest proximity as the strongest predictor of building collisions (99.7% and 96.5% variable contribution, respectively; Table 1 and Appendix S3). Similarly, our models suggest Asian Emerald Dove collisions are strongly linked to forest proximity at both the fine and landscape scale (48.5% and 51.5% variable contribution, respectively, Table 1 and Appendix S3), although the fine scale model also identifies low building height as a predictor (45.3%) while the landscape-scale model identifies building density (43.9%).

Plotting the predicted collision distribution grids for each taxon highlights striking differences in the expected collision landscapes (Fig. 4). Although the overall landscape-wide collision risk is high across all modelled taxa, for both pittas and bitterns, collision hotspots appear to be concentrated around the Central Business District and downtown commercial district where building density and blue light pollution levels are high (Figs. 4C and 4D), although the bitterns also show elevated collision risk in industrial areas in the west. In contrast, the *Ficedula* flycatchers, Pink-necked Green Pigeon, and Asian Emerald Dove are more likely to collide with buildings in areas close to wooded cover, such as in areas fringing the Central Nature Reserve and the Southern Ridges (Figs. 4A, 4B, and 4E).

### Future Collision Landscapes

Projecting the best-fit models onto future land-use scenarios, we find that future urban developments are likely to pose some additional collision risk to birds strongly affected by forest proximity, owing to the encroachment of these developments into the edges of wooded areas (Fig. 5). More importantly, an expected future increase in blue light pollution is likely to dramatically increase the landscape-wide collision risk for migratory species. For Blue-winged Pittas in particular, scenarios with blue light pollution levels exceeding 40% of present-day streetlight intensity are projected to increase the mean landscape-wide collision risk by at least 10.1% (Fig. 6 and Appendix S8).

**Figure 6:**
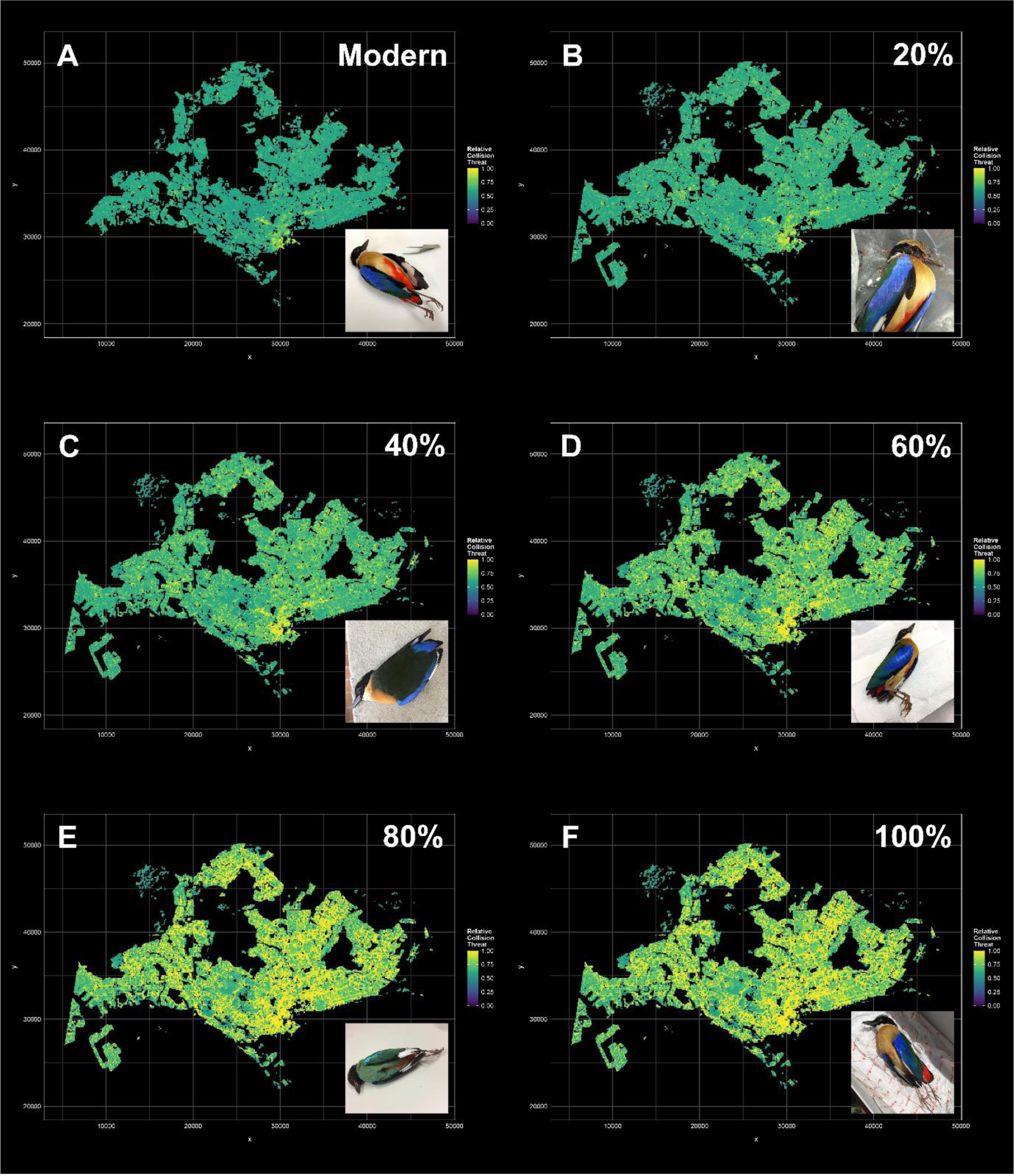
Forecast collision risk maps for the pittas (*Pitta sp.*) for five projected fine-scale scenarios of future nocturnal blue light pollution resulting from Singapore’s transition to white LED streetlights from red/orange sodium vapor lamps. These scenarios correspond with (A) 0% (i.e. modern-day), (B) 20%, (C) 40%, (D) 60%, (E) 80%, and (F) 100% increases in blue light output relative to present-day levels of red streetlight pollution and suggest that a >40% increase in blue light pollution is likely to lead to major increases in landscape-wide relative collision risk.

As for the suitability of our model projections, our MESS analyses indicate no major differences between the training and prediction datasets, indicating that our projections do not require significant extrapolation beyond the parameter space of the training dataset (Appendices S9-S16). It is worth noting, however, that models incorporating blue light pollution as a predictor variable tend to exhibit relatively lower MESS suitabilities, especially for areas with already high levels of blue light pollution and for models in which the projected increase in blue light pollution is high. The relative effect of these deviations is unlikely to affect our projections, however, since the unsuitable area is small relative to the overall projected area.

## Discussion

### MaxEnt is a suitable method for modeling bird-building collisions

In addition to being one of the first multivariate analyses of bird-building collisions from Asia, our study is also one of the first to apply maximum entropy modeling techniques to studying the drivers of bird-building collisions at a landscape scale. Our results demonstrate that MaxEnt can produce robust model outputs with biologically meaningful inferences providing baseline assessments of the collision risk landscape, even with opportunistically sampled community science data. Collision risk maps such as ours (Figs. 4-6) can be used to prioritize high collision risk areas for targeted mitigation efforts. Furthermore, the ability to project MaxEnt models onto future development scenarios makes this tool potentially useful for urban planners and policymakers to facilitate the incorporation of preemptive collision mitigation measures into development plans.

### Drivers of bird-building collisions vary among taxa

Similar to studies from North America (Loss et al., 2019; Elmore et al.,, 2020), we find that the main drivers of bird-building collisions vary among taxa. In contrast to those previous studies, however, we observe that building size is a relatively poor predictor of bird-building collisions, and instead find that building density (i.e. the overall space occupied by buildings per pixel), blue light pollution, forest proximity, and building height are better predictors of collision risk. It is unclear why building size emerged as a relatively good predictor in other studies. It is possible that this may be a function of spatial scale, since when contrasted against 100,000 buildings across an entire city, individual large buildings are less likely to be major obstacles to bird movement relative to dense clusters of buildings. Similarly, at very fine spatial scales such as university campuses and city centers, where building densities are relatively high and homogeneous, the relative effect of individual large buildings should be comparatively stronger.

### Increased Blue Light Pollution Exacerbates Migratory Bird Collision Risk

One of the more surprising results of this study was the fact that blue light pollution is significantly associated with migratory bird-building collisions, particularly in pittas. While the association between light pollution and increased collision risk is well-documented (Lao et al., 2020; Van Doren et al., 2021), relatively few studies have explored the impact of different light spectra. And while pittas are known to be strongly attracted to bright lights during migration, as evidenced by historical records of individuals captured at One Fathom Bank lighthouse in the Straits of Malacca (Robinson & Boden Kloss, 1922) and at the MAPS banding station at Fraser’s Hill, Malaysia (Wells, 1992), no specific attraction to blue light has been documented in this taxon to date. Our finding that blue, but not red, light pollution is associated with increased collision risk among migratory birds echoes a recent study by Zhao et al. (2020), who report elevated catch rates of migratory birds at mist nets illuminated by blue light at night in Yunnan, China, and suggest that future studies need to account for the differential effects of light color temperature on nocturnal migrants. The finding that migrants such as pittas appear acutely sensitive to blue light pollution is particularly concerning given recent changes to Singapore’s light pollution profile. While the bulk of blue light pollution in Singapore was restricted to downtown areas and shopping districts during our study period, the Singapore government has since 2022 replaced most of the country’s sodium-vapor streetlights (which emit orange/red light) with LEDs (Abdullah, 2017), which would dramatically increase island-wide levels of blue light pollution. Our forecasting analyses indicate that increases in streetlamp-induced blue light pollution levels greater than or equal to 40% of present-day red light levels are likely to lead to significant increases in island-wide collision risk for migratory pittas and potentially other migratory species (Fig. 6).

A concerted effort by conservationists is thus needed to reduce the impacts of the LED transition on migratory birds, not only in Singapore, but also in other cities located along migratory pathways. The LED transition underway in many developed countries is part of global efforts to reduce energy consumption and address climate change and is unlikely to be abandoned because of its impact on migrant birds. As such, solutions aimed at mitigating the negative effects of blue light on migratory birds should focus on attenuating the amount of blue light emitted by LED lights and other light sources during peak migratory months, such as by permanently or temporarily deploying LED streetlights with a lower color temperature (∼2700 – 3000 K), which emit warmer, more red-shifted light, instead of LEDs with a higher color temperature (∼4000 – 5000 K), which emit cooler light with a higher blue fraction. Other solutions may be architectural or design-related in nature, such as addressing the way light is directionally shielded, thereby minimizing the exposure of blue light to overflying birds. Further research is also needed into the physiological effects of blue light on night-migrating birds and whether these effects are uniform across migratory avian taxa.

### Forest Edges Are Collision Hotspots

Although forest proximity has not been previously identified as a driver of bird-building collisions, it is likely that the importance of this predictor is partly explained by higher densities of forest-dwelling species near forest edges relative to elsewhere in the urban matrix (Sabo et al., 2016; Brisque et al., 2017), which is known to correlate with increased chances of building collisions (Hager et al., 2008). At the same time, collision rates are likely exacerbated by the fact that forest-dwelling frugivores such as green pigeons and emerald doves are highly dispersive and move between forest patches to forage (Cros et al., 2020), thereby encountering buildings near the forest edge. It is unclear, however, why other edge-dwelling frugivorous species such as bulbuls or barbets appear to be less affected by bird-building collisions, especially since barbets are strongly overrepresented in building collision reports in other East Asian localities such as Taiwan (Wang Ling-Min, pers. comm.). Forest proximity also emerged as a driver of collision risk in some migratory taxa such as Yellow-rumped Flycatchers, but not in others such as pittas. Yellow-rumped Flycatchers are known to migrate during the night and seek out forest patches similar to their breeding habitat during the day to forage and rest on migration, putting them at greater risk of collision with buildings in forest-adjacent areas. Our finding that forest proximity is a strong predictor of collisions in both migratory and non-migratory taxa suggests that buildings near forest habitats are hotspots of bird collisions, and are therefore sites most in need of collision mitigation measures.

To a lesser extent, we also observed that vegetation density is broadly correlated with collision risk for migratory and non-migratory species, especially at coarser spatial scales, although we did not observe any correlation between vegetation density and any of the ‘supercollider’ species. While vegetation density may play a role in attracting both migratory and non-migratory birds at coarse scales, at finer spatial and taxonomic scales, vegetation quality likely matters more in determining the risk of building collisions. Alternatively, our results could also point to the limitations of the NDVI metric as a measure of vegetation density, and alternative metrics such as the Leaf Area Index (LAI) or Enhanced Vegetation Index (EVI) should be considered in future work.

Our research highlights how the drivers of bird-building collisions may differ between temperate and tropical latitudes. The relative abundance of non-migratory forest dwelling frugivores such as green pigeons and emerald doves in tropical forests, combined with the elevated susceptibility of tropical pigeons to building collisions (Ocampo-Penuela et al., 2016; Santos et al., 2017), likely contribute to higher rates of non-migratory bird-building collisions relative to temperate latitudes, and thus the concomitant importance of forest proximity as a key driver of collision risk. More importantly, our results add to the long list of negative impacts of forest fragmentation, since fragmented forest landscapes are characterized by more edges with high collision frequencies.

### Challenges and Opportunities

Despite our dataset spanning seven years, the difficulty of diagnosing cause of death, the relatively high carcass disposal rates across Singapore (Tan et al., 2017), as well as the highly heterogeneous detection rates of community scientists make large sample sizes difficult to achieve. Although MaxEnt may be a suitable method for analyzing opportunistically obtained bird-building collision data, some of our models performed poorly, especially in the case of the Black-backed Kingfisher, likely pointing to the effect of small sample sizes. Combining community science with more systematic survey efforts may help address the sample size limitation so long as differences in survey effort are accounted for.

Our models benefited from the availability of highly detailed fine-scale and landscape-scale open-source building data, as well as high-fidelity land-use forecasts, which may remain elusive in many other countries. In the future, other predictor variables that have been strongly associated with bird-building collisions, such as façade glass cover and the amount of light reflected off glass façades, could be included in similar research (Hager et al., 2017, Lao et al., 2020). At present, these variables are challenging to quantify at the city-wide scale. Future analyses, however, should be able to address this limitation by combining emerging citywide 3D photogrammetric models with machine learning and computer vision methods to estimate fine scale urban characteristics such as glass area across entire cities.

Generating environmental predictor layers can be challenging and time-consuming to prepare. For example, generating the raster for building height in this study entailed manually annotating height data from individual buildings in the OneMap3D portal onto our shapefile of building polygons, which required approximately two months of continuous work to complete. With increasing data availability, it is hoped that the incorporation of such data will be more straightforward in the future.

The analytical approach introduced in this study can serve as a blueprint for more rigorous and applied analyses of bird-building collisions in cities worldwide. Such analyses can subsequently be translated into policies aimed at reducing the incidence of bird-building collisions in urban areas.

## Acknowledgements

We thank the hundreds of community scientists from across Singapore who have contributed bird mortality records to this project.

## Supporting Information

Additional supporting information may be found in the online version of the article at the publisher’s website.

## References

Abdullah, Z. (2017). LTA installing smarter, energy-saving street lights. The Straits Times. Available from https://www.straitstimes.com/singapore/lta-installing-smarter-energy-saving-street-lights.

Barton, C.M., Riding, C.S., & Loss, S.R. (2017). Magnitude and correlates of bird collisions at glass bus shelters in an urban landscape. PLOS ONE 12(6), 12:e0178667.

Basilio, L.G., Moreno, D.J., & Piratelli, A.J. (2020). Main causes of bird-window collisions: a review. Anais da Academia Brasileira de Ciências 92(1).

Bayne, E.M., Scobie, C.A., & Rawson-Clark, M. (2012). Factors influencing the annual risk of bird–window collisions at residential structures in Alberta, Canada. Wildlife Research 39:583.

Biljecki, F. (2020). Exploration of open data in Southeast Asia to generate 3D building models. ISPRS Annals of Photogrammetry, Remote Sensing and Spatial Information Sciences VI-4/W1–2020:37–44.

Borden, C.W., Lockhart, O.M., Jones, A.W., & Lyons, M.S. (2010). Seasonal, taxonomic, and local habitat components of bird-window collisions on an urban university campus in Cleveland, OH. Ohio Journal of Science 110:44–52.

Brisque, T., Campos-Silva, L.A., & Piratelli, A.J. (2017). Relationship between bird-of-prey decals and bird-window collisions on a Brazilian university campus. Zoologia 34:1–8.

Cobos, M.E., Peterson, A.T., Barve, N., & Osorio-Olvera, L. (2019). kuenm: an R package for detailed development of ecological niche models using Maxent. PeerJ, 7:e6281.

Cros, E., Ng, E.Y.X., Oh, R.R.Y., Tang, Q., Benedick, S., Edwards, D.P., Tomassi, S., Irestedt, M., Ericson, P.G.P., & Rheindt, F.E. (2020). Fine-scale barriers to connectivity across a fragmented South-East Asian landscape in six songbird species. Evolutionary Applications 13:1026–1036.

Cusa, M., Jackson, D.A., & Mesure, M. (2015). Window collisions by migratory bird species: urban geographical patterns and habitat associations. Urban Ecosystems 18:1427–1446.

Delignette-Muller, M.L., & Dutang, C. (2015). fitdistrplus: An R Package for Fitting Distributions. Journal of Statistical Software 64(4):1–34.

Elith, J., Phillips, S.J., Hastie, T., Dudík, M., Chee, Y.E., & Yates, C.J. (2010). A statistical explanation of MaxEnt for ecologists. Diversity and Distributions 17:43–57.

Elmore, J. A., Hager, S. B., Cosentino, B. J., O’Connell, T. J., Riding, C. S., Anderson, M. L., Bakermans, M. H., Boves, T. J., Brandes, D., Butler, E. M., Butler, M. W., Cagle, N. L., Calderón-Parra, R., Capparella, A. P., Chen, A., Cipollini, K., Conkey, A. A. T., Contreras, T. A., Cooper, R. I., … Loss, S. R. (2020). Correlates of bird collisions with buildings across three North American countries. Conservation Biology 35(2): 654–665.

Freymueller, N.F. (2020). Niche dynamics of the felid guild following the Pleistocene Megafaunal Extinction (Unpublished Master’s thesis). University of New Mexico, Albuquerque, USA.

Gaw, L.Y.F., Yee, A.T.K., & Richards D.R. (2019). A High-Resolution Map of Singapore’s Terrestrial Ecosystems. Data 4:116.

Gelb, Y., & Delacretaz, N. (2009). Windows and Vegetation: Primary Factors in Manhattan Bird Collisions. Northeastern Naturalist 16:455–470.

Gomes, L., Grilo, C., & Mira, A. (2008). Identification methods and deterministic factors of owl roadkill hotspot locations in Mediterranean landscapes. Ecological Research 24:355–370.

Gómez-Martínez, M.A., Klem, D. Jr., Rojas-Soto, O., González-García, F., & MacGregor-Fors, I. (2019). Window strikes: bird collisions in a Neotropical green city. Urban Ecosystems 22:699–708.

Hager, S.B., Trudell, H., McKay, K.J., Crandall, S.M., & Mayer, L. (2008). Bird density and mortality at windows. The Wilson Journal of Ornithology 120:550–564.

Hager, S. B., Cosentino, B. J., Aguilar-Gómez, M. A., Anderson, M. L., Bakermans, M., Boves, T. J., Brandes, D., Butler, M. W., Butler, E. M., Cagle, N. L., Calderón-Parra, R., Capparella, A. P., Chen, A., Cipollini, K., Conkey, A. A. T., Contreras, T. A., Cooper, R. I., Corbin, C. E., Curry, R. L., … Zuria, I. (2017). Continent-wide analysis of how urbanization affects bird-window collision mortality in North America. Biological Conservation 212:209–215.

Hijmans, R.J., Phillips, S., Leathwick, J., & Elith, J. (2015). Dismo: Species Distribution Modeling. R Package Version 1.0–12. Available from http://CRAN.R-project.org/package=dismo.

Klem, D. (1989). Bird-Window Collisions. The Wilson Bulletin 101: 606–620.

Land Transport Authority of Singapore. (2017). Smarter and More Energy-Efficient Street Lighting System by 2022. Available from: https://www.lta.gov.sg/content/ltagov/en/newsroom/2017/1/2/smarter-and-more-energy-efficient-street-lighting-system-by-2022.html

Lao, S., Robertson, B.A., Anderson, A.W., Blair, R.B., Eckles, J.W., Turner, R.J., & Loss, S.R. (2020). The influence of artificial light at night and polarized light on bird-building collisions. Biological Conservation 241.

Longcore, T., & Rich, C. (2004). Ecological light pollution. Frontiers in Ecology and the Environment 2:191–198.

Loss, S.R., Will, T., & Marra, P.P. (2013). The impact of free-ranging domestic cats on wildlife of the United States. Nature Communications 4.

Loss, S.R., Will, T., Loss, S.S., & Marra, P.P. (2014). Bird–building collisions in the United States: Estimates of annual mortality and species vulnerability. The Condor 116:8–23.

Loss, S.R., Will, T., & Marra, P.P. (2015). Direct Mortality of Birds from Anthropogenic Causes. Annual Review of Ecology, Evolution, and Systematics 46:99–120.

Low, B.W., Yong, D.L., Tan, D.J.X., Owyong, A., & Chia, A. (2017). Migratory bird collisions with man-made structures in South-East Asia: a case study from Singapore. BirdingASIA 27:107–111

Menacho-Odio, R.M., Garro-Cruz, M., & Arévalo, J.E. (2019). Ecology, endemism, and conservation status of birds that collide with glass windows in Monteverde, Costa Rica. Revista de Biología Tropical, 67(2) Suplemento, S326-S345.

Muscarella, R., Galante, P.J., Soley-Guardia, M., Boria, R.A., Kass, J.M., Uriarte, M., & Anderson, R.P. (2014). ENMeval: An R package for conducting spatially independent evaluations and estimating optimal model complexity for Maxent ecological niche models. Methods in Ecology and Evolution 5:1198–1205.

Ocampo-Peñuela, N., Peñuela-Recio, L., & Ocampo-Durán, Á. (2015). Decals prevent bird-window collisions at residences: a successful case study from Colombia. Ornitología Colombiana, 15:84–91.

Ocampo-Peñuela, N., Winton, R.S., Wu, C.J., Zambello, E., Wittig, T.W., & Cagle, N.L. (2016). Patterns of bird-window collisions inform mitigation on a university campus. PeerJ 4:e1652.

Pearson, R.G., Raxworthy, C.J., Nakamura, M., Townsend Peterson, A. (2006). Predicting species distributions from small numbers of occurrence records: a test case using cryptic geckos in Madagascar. Journal of Biogeography 34: 102–117.

Peterson, A.T., Papeş, M., & Soberón, J. (2008). Rethinking receiver operating characteristic analysis applications in ecological niche modeling. Ecological Modeling 213(1):63–72.

Peterson, A.T., & Samy, A.M. (2016). Geographic potential of disease caused by Ebola and Marburg viruses in Africa. Acta Tropica 162:114–124.

Phillips, S.J., Anderson, R.P., & Schapire, R.E. (2006). Maximum entropy modeling of species geographic distributions. Ecological Modeling 190:231–259.

Phillips, S.J., Anderson, R.P., Dudik, M., Schapire, R.E., & Blair, M.E. (2017). Opening the black box: an open-source release of Maxent. Ecography 40(7):887–893.

Rebolo-Ifrán, N., di Virgilio, A., & Lambertucci, S.A. (2019). Drivers of bird-window collisions in southern South America: a two-scale assessment applying citizen science. Scientific Reports 9.

Robinson, H.C., & Boden Kloss, C. (1922). Birds from the One Fathom Bank Lighthouse, Straits of Malacca. Journal of the Federated Malay Straits Museum 10:253–255.

Sabo, A.M., Hagemeyer, N.D.G., Lahey, A.S., & Walters, E.L. (2016). Local avian density influences risk of mortality from window strikes. PeerJ 4:e2170.

Santos, L.P.S., de Abreu, V.F., & de Vasconcelos, M.F. (2017). Bird mortality due to collisions in glass panes on an Important Bird Area of southeastern Brazil. Revista Brasileira de Ornitologia 25:90–101.

Smeraldo, S., Bosso, L., Fraissinet, M., Bordignon, L., Brunelli, M., Ancillotto, L., & Russo, D. (2020). Modeling risks posed by wind turbines and power lines to soaring birds: the black stork (*Ciconia nigra*) in Italy as a case study. Biodiversity and Conservation 29:1959–1976.

Tan, D.J.X., Yong, D.L., Low, B.W., Owyong, A., & Chia, A. (2017). Anthropogenic sources of non-migratory avian mortalities in Singapore. International Journal of Tropical Veterinary and Biomedical Research 2: 17–27.

Van Doren, B.M., Willard, D.E., Hennen, M., Horton, K.G., Stuber, E.F., Sheldon, D., Sivakumar, A.H., Wang, J., Farnsworth, A., & Winger, B.M. (2021). Drivers of fatal bird collisions in an urban center. Proceedings of the National Academy of Sciences 118 (24).

Warren, D.L., & Seifert, S.N. (2011). Ecological niche modeling in Maxent: The importance of model complexity and the performance of model selection criteria. Ecological Applications 25:335–342.

Wells, D.R. (1992). Night Migration at Fraser’s Hill. Bulletin of the Oriental Bird Club 16:21–25.

Wickham H (2016). ggplot2: Elegant Graphics for Data Analysis. Springer-Verlag New York. ISBN 978-3-319-24277-4, https://ggplot2.tidyverse.org.

Winger, B.M., Weeks, B.C., Farnsworth, A., Jones, A.W., Hennen, M., & Willard, D.E. (2019). Nocturnal flight-calling behaviour predicts vulnerability to artificial light in migratory birds. Proceedings of the Royal Society B: Biological Sciences 286:20190364.

Zhao, X., Zhang, M., Che, X., & Zou, F. (2020). Blue light attracts nocturnally migrating birds. The Condor 122.

