## Supporting Information for "Disentangling the biotic and abiotic drivers of bird-building collisions in a tropical Asian city using ecological niche modeling"

Appendix S1: Best-fit gamma distribution model parameters for each predictor variable and future land-use type category (per the 2019 Singapore Master Plan). These distributions were fit based on baseline values extracted from present-day maps, and the best-fit parameters were used to generate five randomised rasters for each land-use type and predictor variable to model the likely future landscape of Singapore based on the 2019 Singapore Master Plan.

| Predictor | Landuse | Mean | alpha | rate | lambda | Remarks |
| --- | --- | --- | --- | --- | --- | --- |
| BuildingArea_Mean | Business1 | 5087.61534 | 1.04151832 | 0.00204787 | 488.311769 | Scaled by 1/10 |
| BuildingArea_Mean | Business2 | 6934.92743 | 0.86949093 | 0.01253502 | 79.7764982 | Scaled by 1/100 |
| BuildingArea_Mean | BusinessPark | 9160.12174 | 0.76326652 | 0.00833122 | 120.030512 | Scaled by 1/100 |
| BuildingArea_Mean | Civic | 4444.584 | 0.75725894 | 0.01703502 | 58.7026021 | Scaled by 1/100 |
| BuildingArea_Mean | Commercial | 6647.799 | 0.62923397 | 0.009464 | 105.663523 | Scaled by 1/100 |
| BuildingArea_Mean | Educational | 4546.666 | 0.7711546 | 0.01695589 | 58.9765562 | Scaled by 1/100 |
| BuildingArea_Mean | Medical | 5663.23 | 0.67134486 | 0.01184935 | 84.3928148 | Scaled by 1/100 |
| BuildingArea_Mean | Residential | 2202.919 | 1.38977974 | 0.06309303 | 15.8496113 | Scaled by 1/100 |
| BuildingArea_Mean | Sport | 5627.51 | 0.40884232 | 0.00726677 | 137.612798 | Scaled by 1/100 |
| BuildingArea_Mean | Transport | 7507.271 | 0.57957704 | 0.00771895 | 129.551366 | Scaled by 1/100 |
| BuildingArea_Mean | Utilities | 3359.15 | 0.7403119 | 0.02205157 | 45.348245 | Scaled by 1/100 |
| RedLight | Business1 | 84.195753 | 5.23156309 | 0.06214301 | 16.0919144 | Unscaled |
| RedLight | Business2 | 69.905822 | 3.32417015 | 0.04755321 | 21.0290746 | Unscaled |
| RedLight | BusinessPark | 80.41449 | 4.4353353 | 0.05515216 | 18.1316561 | Unscaled |
| RedLight | Civic | 83.277688 | 3.70199086 | 0.04445174 | 22.4963072 | Unscaled |
| RedLight | Commercial | 98.2880355 | 5.95302638 | 0.06056455 | 16.511309 | Unscaled |
| RedLight | Educational | 76.080144 | 4.90226553 | 0.06443098 | 15.5204841 | Unscaled |
| RedLight | Medical | 89.83725 | 5.93046201 | 0.06600883 | 15.1494883 | Unscaled |
| RedLight | Residential | 82.356233 | 5.57321481 | 0.06767322 | 14.776894 | Unscaled |
| RedLight | Sport | 42.0763389 | 1.3783666 | 0.03276291 | 30.5223193 | Unscaled |
| RedLight | Transport | 88.985991 | 4.10653578 | 0.04614886 | 21.6690076 | Unscaled |
| RedLight | Utilities | 64.943929 | 1.87883179 | 0.02892723 | 34.5695042 | Unscaled |
| BuildingFootprint | Business1 | 2804.782 | 1.46381511 | 0.00521956 | 191.587177 | Scaled by 1/10 |
| BuildingFootprint | Business2 | 2927.277 | 1.09518695 | 0.00373904 | 267.448615 | Scaled by 1/10 |
| BuildingFootprint | BusinessPark | 2694.665 | 1.19697369 | 0.00444428 | 225.008477 | Scaled by 1/10 |
| BuildingFootprint | Civic | 2556.481 | 1.37606096 | 0.00538465 | 185.713195 | Scaled by 1/10 |
| BuildingFootprint | Commercial | 2858.286 | 1.47567386 | 0.00516245 | 193.706552 | Scaled by 1/10 |
| BuildingFootprint | Educational | 2441.236 | 1.75143555 | 0.0071751 | 139.37086 | Scaled by 1/10 |
| BuildingFootprint | Medical | 2542.78 | 1.82165703 | 0.00716222 | 139.621436 | Scaled by 1/10 |
| BuildingFootprint | Residential | 2289.025 | 1.74405726 | 0.00761784 | 131.270875 | Scaled by 1/10 |
| BuildingFootprint | Sport | 1414.302 | 0.7177994 | 0.0050758 | 197.013318 | Scaled by 1/10 |
| BuildingFootprint | Transport | 2499.129 | 1.06949865 | 0.00427874 | 233.713827 | Scaled by 1/10 |
| BuildingFootprint | Utilities | 2356.773 | 0.98003535 | 0.00415828 | 240.484335 | Scaled by 1/10 |
| BuildingFootprint_mean | Business1 | 1194.917 | 1.16972701 | 0.00978791 | 102.166909 | Scaled by 1/10 |
| BuildingFootprint_mean | Business2 | 1436.304 | 0.95573 | 0.00665387 | 150.288501 | Scaled by 1/10 |
| BuildingFootprint_mean | BusinessPark | 1454.3 | 1.08063661 | 0.00742963 | 134.596205 | Scaled by 1/10 |
| BuildingFootprint_mean | Civic | 956.1461 | 1.09617893 | 0.01146079 | 87.25402 | Scaled by 1/10 |
| BuildingFootprint_mean | Commercial | 1098.817 | 1.08341779 | 0.00986212 | 101.398128 | Scaled by 1/10 |
| BuildingFootprint_mean | Educational | 946.305 | 1.2270614 | 0.01296702 | 77.1187212 | Scaled by 1/10 |
| BuildingFootprint_mean | Medical | 1031.933 | 1.14263204 | 0.01106955 | 90.3379089 | Scaled by 1/10 |
| BuildingFootprint_mean | Residential | 694.8362 | 1.58264642 | 0.02277516 | 43.9074852 | Scaled by 1/10 |
| BuildingFootprint_mean | Sport | 723.1909 | 0.72805521 | 0.01006529 | 99.3513351 | Scaled by 1/10 |
| BuildingFootprint_mean | Transport | 1180.614 | 0.82423917 | 0.00698099 | 143.246138 | Scaled by 1/10 |
| BuildingFootprint_mean | Utilities | 894.5729 | 0.85777077 | 0.00958917 | 104.284291 | Scaled by 1/10 |
| BuildingPerimeter | Business1 | 276.5495 | 2.03404961 | 0.00735581 | 135.947033 | Unscaled |
| BuildingPerimeter | Business2 | 222.4922 | 1.84749671 | 0.00830449 | 120.416859 | Unscaled |
| BuildingPerimeter | BusinessPark | 209.625 | 1.79493959 | 0.00856179 | 116.797992 | Unscaled |
| BuildingPerimeter | Civic | 319.1613 | 1.75250605 | 0.00549115 | 182.111381 | Unscaled |
| BuildingPerimeter | Commercial | 329.6999 | 1.94500808 | 0.00589984 | 169.496151 | Unscaled |
| BuildingPerimeter | Educational | 361.8441 | 2.07426683 | 0.00573276 | 174.436079 | Unscaled |
| BuildingPerimeter | Medical | 340.3345 | 2.22661969 | 0.00654212 | 152.855766 | Unscaled |
| BuildingPerimeter | Residential | 370.045 | 2.17585197 | 0.00588287 | 169.985174 | Unscaled |
| BuildingPerimeter | Sport | 191.7444 | 1.13126161 | 0.0058994 | 169.508649 | Unscaled |
| BuildingPerimeter | Transport | 280.05 | 1.34218489 | 0.00479332 | 208.623799 | Unscaled |
| BuildingPerimeter | Utilities | 295.917 | 1.22134266 | 0.00412787 | 242.255456 | Unscaled |
| NDVI | Business1 | 0.2026484 | 94.02157 | 78.17845 | 0.01279125 | Scaled by NDVI +1 |
| NDVI | Business2 | 0.146025 | 93.22736 | 81.34823 | 0.01229283 | Scaled by NDVI +1 |
| NDVI | BusinessPark | 0.2276204 | 91.44567 | 74.49004 | 0.01342461 | Scaled by NDVI +1 |
| NDVI | Civic | 0.2102632 | 84.00827 | 69.41242 | 0.01440664 | Scaled by NDVI +1 |
| NDVI | Commercial | 0.1808643 | 108.1556 | 91.5905 | 0.01091816 | Scaled by NDVI +1 |
| NDVI | Educational | 0.2192333 | 122.4373 | 100.4205 | 0.00995813 | Scaled by NDVI +1 |
| NDVI | Medical | 0.2052903 | 123.698 | 102.6284 | 0.00974389 | Scaled by NDVI +1 |
| NDVI | Residential | 0.2129978 | 123.3972 | 101.7288 | 0.00983006 | Scaled by NDVI +1 |
| NDVI | Sport | 0.327513 | 57.92148 | 43.63181 | 0.02291906 | Scaled by NDVI +1 |
| NDVI | Transport | 0.1993078 | 97.45129 | 81.25672 | 0.01230667 | Scaled by NDVI +1 |
| NDVI | Utilities | 0.230698 | 70.79244 | 57.52189 | 0.01738469 | Scaled by NDVI +1 |
| BlueLight_20 | Business1 | 41.60791 | 3.76527547 | 0.09048657 | 11.0513638 | Unscaled |
| BlueLight_20 | Business2 | 31.22508 | 2.72840207 | 0.08737723 | 11.4446292 | Unscaled |
| BlueLight_20 | BusinessPark | 39.65406 | 2.9198664 | 0.0736334 | 13.5807935 | Unscaled |
| BlueLight_20 | Civic | 40.86883 | 2.55001087 | 0.06239983 | 16.0256847 | Unscaled |
| BlueLight_20 | Commercial | 53.13692 | 3.27947353 | 0.06170965 | 16.2049209 | Unscaled |
| BlueLight_20 | Educational | 35.32207 | 3.32280984 | 0.09407619 | 10.6296822 | Unscaled |
| BlueLight_20 | Medical | 43.44751 | 3.74904637 | 0.08628245 | 11.5898424 | Unscaled |
| BlueLight_20 | Residential | 39.5152 | 3.8288821 | 0.0969003 | 10.3198855 | Unscaled |
| BlueLight_20 | Sport | 20.31758 | 1.57008926 | 0.07728054 | 12.9398682 | Unscaled |
| BlueLight_20 | Transport | 40.78246 | 3.0946238 | 0.07589963 | 13.1752948 | Unscaled |
| BlueLight_20 | Utilities | 29.30703 | 1.8601871 | 0.0634626 | 15.7573122 | Unscaled |
| BlueLight_40 | Business1 | 56.13173 | 3.2915395 | 0.05863813 | 17.0537498 | Unscaled |
| BlueLight_40 | Business2 | 39.73397 | 2.1919719 | 0.05515937 | 18.1292861 | Unscaled |
| BlueLight_40 | BusinessPark | 53.43841 | 2.69777137 | 0.05047613 | 19.8113445 | Unscaled |
| BlueLight_40 | Civic | 53.99416 | 2.18133342 | 0.04039499 | 24.7555452 | Unscaled |
| BlueLight_40 | Commercial | 69.65007 | 2.99363966 | 0.04298119 | 23.2659915 | Unscaled |
| BlueLight_40 | Educational | 46.86777 | 2.66891891 | 0.05693981 | 17.5624049 | Unscaled |
| BlueLight_40 | Medical | 58.31731 | 3.24546958 | 0.05565661 | 17.9673178 | Unscaled |
| BlueLight_40 | Residential | 52.65392 | 3.1304648 | 0.05945968 | 16.8181194 | Unscaled |
| BlueLight_40 | Sport | 25.75139 | 1.22853514 | 0.04771567 | 20.9574758 | Unscaled |
| BlueLight_40 | Transport | 54.09776 | 2.6150262 | 0.0483386 | 20.687401 | Unscaled |
| BlueLight_40 | Utilities | 38.82227 | 1.52473706 | 0.03928073 | 25.4577754 | Unscaled |
| BlueLight_60 | Business1 | 70.65555 | 2.88077623 | 0.04077367 | 24.5256314 | Unscaled |
| BlueLight_60 | Business2 | 48.24286 | 1.82685825 | 0.03788038 | 26.3988904 | Unscaled |
| BlueLight_60 | BusinessPark | 67.22276 | 2.45149534 | 0.03647799 | 27.4137912 | Unscaled |
| BlueLight_60 | Civic | 67.1195 | 1.89997294 | 0.02830821 | 35.3254409 | Unscaled |
| BlueLight_60 | Commercial | 86.16322 | 2.67850085 | 0.03108499 | 32.1698672 | Unscaled |
| BlueLight_60 | Educational | 58.41347 | 2.22925967 | 0.03816289 | 26.2034662 | Unscaled |
| BlueLight_60 | Medical | 73.18711 | 2.8132292 | 0.0384417 | 26.0134177 | Unscaled |
| BlueLight_60 | Residential | 65.79263 | 2.63009277 | 0.03997293 | 25.0169302 | Unscaled |
| BlueLight_60 | Sport | 31.1852 | 1.03454082 | 0.03317792 | 30.1405272 | Unscaled |
| BlueLight_60 | Transport | 67.41306 | 2.231995 | 0.0331069 | 30.2051838 | Unscaled |
| BlueLight_60 | Utilities | 48.33751 | 1.3142823 | 0.02718541 | 36.7844369 | Unscaled |
| BlueLight_80 | Business1 | 85.17937 | 2.58152036 | 0.03030871 | 32.993816 | Unscaled |
| BlueLight_80 | Business2 | 56.75174 | 1.58411183 | 0.02790711 | 35.8331622 | Unscaled |
| BlueLight_80 | BusinessPark | 81.0071 | 2.24954035 | 0.02776759 | 36.0132082 | Unscaled |
| BlueLight_80 | Civic | 80.24484 | 1.70042461 | 0.02119234 | 47.1868609 | Unscaled |
| BlueLight_80 | Commercial | 102.6764 | 2.42300144 | 0.02359876 | 42.3751078 | Unscaled |
| BlueLight_80 | Educational | 69.95917 | 1.93954599 | 0.02772111 | 36.0735916 | Unscaled |
| BlueLight_80 | Medical | 88.05692 | 2.49824882 | 0.02836752 | 35.2515835 | Unscaled |
| BlueLight_80 | Residential | 78.93135 | 2.29320051 | 0.02905538 | 34.417034 | Unscaled |
| BlueLight_80 | Sport | 36.61902 | 0.9109526 | 0.02487179 | 40.2061934 | Unscaled |
| BlueLight_80 | Transport | 80.72836 | 1.96323243 | 0.02431749 | 41.1226652 | Unscaled |
| BlueLight_80 | Utilities | 57.85275 | 1.17357159 | 0.02028313 | 49.3020555 | Unscaled |
| BlueLight_100 | Business1 | 99.70319 | 2.35748874 | 0.02363985 | 42.3014528 | Unscaled |
| BlueLight_100 | Business2 | 65.26063 | 1.413711 | 0.0216569 | 46.1746603 | Unscaled |
| BlueLight_100 | BusinessPark | 94.79145 | 2.0881393 | 0.02202388 | 45.4052601 | Unscaled |
| BlueLight_100 | Civic | 93.37018 | 1.55409456 | 0.01664621 | 60.0737345 | Unscaled |
| BlueLight_100 | Commercial | 119.1895 | 2.22372324 | 0.01866049 | 53.5891608 | Unscaled |
| BlueLight_100 | Educational | 81.50488 | 1.7377079 | 0.0213211 | 46.9018953 | Unscaled |
| BlueLight_100 | Medical | 102.9267 | 2.26660286 | 0.02202401 | 45.4049921 | Unscaled |
| BlueLight_100 | Residential | 92.07006 | 2.05565347 | 0.02232834 | 44.7861328 | Unscaled |
| BlueLight_100 | Sport | 42.05283 | 0.8252784 | 0.0196283 | 50.9468472 | Unscaled |
| BlueLight_100 | Transport | 94.04366 | 1.76979969 | 0.01881849 | 53.1392264 | Unscaled |
| BlueLight_100 | Utilities | 67.368 | 1.07327708 | 0.01593082 | 62.7714079 | Unscaled |
| GreenLight | Business1 | 63.8845192 | 6.169258 | 0.0965753 | 10.3546145 | Unscaled |
| GreenLight | Business2 | 49.35108 | 3.08349469 | 0.06248514 | 16.0038051 | Unscaled |
| GreenLight | BusinessPark | 62.22671 | 4.30349615 | 0.06915659 | 14.4599379 | Unscaled |
| GreenLight | Civic | 61.55093 | 3.63395878 | 0.05904148 | 16.9372448 | Unscaled |
| GreenLight | Commercial | 75.97716 | 6.21407841 | 0.08178265 | 12.2275324 | Unscaled |
| GreenLight | Educational | 51.71992 | 4.41833592 | 0.08541995 | 11.7068671 | Unscaled |
| GreenLight | Medical | 64.19006 | 5.24308872 | 0.08168702 | 12.241847 | Unscaled |
| GreenLight | Residential | 59.41033 | 5.9750426 | 0.1005678 | 9.94354058 | Unscaled |
| GreenLight | Sport | 27.86933 | 1.09412654 | 0.03924745 | 25.4793624 | Unscaled |
| GreenLight | Transport | 72.50522 | 4.73402533 | 0.06528797 | 15.3167574 | Unscaled |
| GreenLight | Utilities | 44.85549 | 1.786471 | 3.982772 | 0.25108141 | Unscaled |
| BuildingHeight_Mean | Business1 | 25.22004 | 3.9616481 | 0.1570926 | 6.36567222 | Unscaled |
| BuildingHeight_Mean | Business2 | 15.74123 | 3.3519167 | 0.2129351 | 4.69626661 | Unscaled |
| BuildingHeight_Mean | BusinessPark | 30.07531 | 4.1932212 | 0.1394335 | 7.17187763 | Unscaled |
| BuildingHeight_Mean | Civic | 21.94739 | 2.9933884 | 0.1364037 | 7.33117943 | Unscaled |
| BuildingHeight_Mean | Commercial | 28.16776 | 1.99255235 | 0.07073671 | 14.1369312 | Unscaled |
| BuildingHeight_Mean | Educational | 24.30604 | 3.9005615 | 0.1604617 | 6.23201674 | Unscaled |
| BuildingHeight_Mean | Medical | 24.54044 | 2.9398504 | 0.1198066 | 8.34678557 | Unscaled |
| BuildingHeight_Mean | Residential | 31.20422 | 3.6547657 | 0.1171201 | 8.53824408 | Unscaled |
| BuildingHeight_Mean | Sport | 14.6999 | 2.1603065 | 0.1469677 | 6.80421616 | Unscaled |
| BuildingHeight_Mean | Transport | 17.26158 | 3.0564194 | 0.1770556 | 5.64794336 | Unscaled |
| BuildingHeight_Mean | Utilities | 15.9916 | 3.224212 | 0.2016031 | 4.96024119 | Unscaled |

Appendix S2: Covariance matrix of fine-scale (100 m x 100 m) and coarse-scale (500 m x 500 m) predictor variables included in this study. Nocturnal green light pollution and total building area were excluded as predictor variables due to high covariance with other predictor variables (covariance > 0.8), but are not shown in this table (due to complete loss of the original raw data files).

|  | BuildingArea_Mean | BuildingArea_Focal | BuildingFootprint_100 | BuildingFootprint_Focal | BuildingHeight_Mean | BuildingHeight_Mean_Focal | BuildingPerimeter | BuildingPerimeter_Focal | ForestProximity | LightPollution_Blue | LightPollution_Blue_Focal | LightPollution_Red | LightPollution_Red_Focal | NDVI | NDVI_Focal |
| --- | --- | --- | --- | --- | --- | --- | --- | --- | --- | --- | --- | --- | --- | --- | --- |
| BuildingArea_Mean | 1 | 0.914446 | 0.10921 | 0.101959 | 0.044325 | 0.005993 | -0.12113 | -0.16166 | 0.039138 | 0.085286 | 0.101216 | 0.184564 | 0.202883 | -0.10728 | -0.15202 |
| BuildingArea_Focal | 0.91445 | 1 | 0.111394 | 0.151533 | 0.041687 | 0.023453 | -0.11238 | -0.15614 | 0.050688 | 0.109105 | 0.139286 | 0.183564 | 0.22018 | -0.10647 | -0.16399 |
| BuildingFootprint_100 | 0.10921 | 0.111394 | 1 | 0.613561 | -0.03038 | 0.055279 | 0.523045 | 0.287434 | 0.169692 | 0.119498 | 0.169827 | 0.003471 | 0.135602 | -0.48329 | -0.35807 |
| BuildingFootprint_Focal | 0.10196 | 0.151533 | 0.613561 | 1 | -0.0219 | 0.130388 | 0.369957 | 0.534605 | 0.306679 | 0.189671 | 0.281538 | 0.146692 | 0.230713 | -0.39137 | -0.57547 |
| BuildingHeight_Mean | 0.04433 | 0.041687 | -0.03038 | -0.0219 | 1 | 0.785513 | -0.07443 | -0.05699 | 0.002246 | 0.272337 | 0.351796 | 0.285638 | 0.353198 | -0.081 | -0.08116 |
| BuildingHeight_Mean_Focal | 0.00599 | 0.023453 | 0.055279 | 0.130388 | 0.785513 | 1 | 0.035958 | 0.096355 | 0.03672 | 0.357932 | 0.4999 | 0.361782 | 0.485248 | -0.12985 | -0.17506 |
| BuildingPerimeter | -0.1211 | -0.11238 | 0.523045 | 0.369957 | -0.07443 | 0.035958 | 1 | 0.741827 | 0.057236 | -0.02615 | -0.02172 | -0.09172 | -0.0052 | -0.27847 | -0.12868 |
| BuildingPerimeter_Focal | -0.1617 | -0.15614 | 0.287434 | 0.534605 | -0.05699 | 0.096355 | 0.741827 | 1 | 0.117943 | -0.02327 | 0.006193 | -0.011268 | 0.040214 | -0.13379 | -0.19559 |
| ForestProximity | 0.03914 | 0.050688 | 0.169692 | 0.306679 | 0.002246 | 0.03672 | 0.057236 | 0.117943 | 1 | 0.137158 | 0.210299 | 0.156035 | 0.22197 | -0.28751 | -0.50565 |
| LightPollution_Blue | 0.08529 | 0.109105 | 0.119498 | 0.189671 | 0.272337 | 0.357932 | -0.02615 | -0.02327 | 0.137158 | 1 | 0.761505 | 0.681263 | 0.529211 | -0.24257 | -0.26428 |
| LightPollution_Blue_Focal | 0.10122 | 0.139286 | 0.169827 | 0.281538 | 0.351796 | 0.4999 | -0.02172 | 0.006193 | 0.210299 | 0.761505 | 1 | 0.570067 | 0.7152 | -0.24325 | -0.36909 |
| LightPollution_Red | 0.18456 | 0.183564 | 0.003471 | 0.146692 | 0.285638 | 0.361782 | -0.09172 | -0.01127 | 0.156035 | 0.681263 | 0.570067 | 1 | 0.801009 | -0.24726 | -0.30493 |
| LightPollution_Red_Focal | 0.20288 | 0.22018 | 0.135602 | 0.230713 | 0.353198 | 0.485248 | -0.0052 | 0.040214 | 0.22197 | 0.529211 | 0.7152 | 0.801009 | 1 | -0.26226 | -0.39487 |
| NDVI | -0.1073 | -0.10647 | -0.48329 | -0.39137 | -0.081 | -0.12985 | -0.27847 | -0.13379 | -0.28751 | -0.24257 | -0.24325 | -0.24726 | -0.26226 | 1 | 0.625997 |
| NDVI_Focal | -0.152 | -0.16399 | -0.35807 | -0.57547 | -0.08116 | -0.17506 | -0.12868 | -0.19559 | -0.50565 | -0.26428 | -0.36909 | -0.304932 | -0.39487 | 0.625997 | 1 |

Appendix S3: Best fit coarse-scale (500 m x 500 m) model loadings for each taxon/migratory phenology, with variable percentage contribution and permutation importance (in parentheses) in the upper table, and model fit metrics reported in the lower table. Taxon names in bold are migratory, while non-bolded taxa are non-migratory. Taxa marked with ‘-’ indicate that the coarse-scale model is not significantly different from the null model (with no predictor variables).

| Variable | ***Pitta*** | ***Ficedula*** | ***Ixobrychus*** | **All Migrants** | *Treron* | *Chalcophaps* | All Residents |
| --- | --- | --- | --- | --- | --- | --- | --- |
| Mean Building Area | 1.97 (1.82) | - | - | 1.21 (1.60) | 0 | 1.01 (0.0318) | 0.310 (3.31) |
| Building Perimeter | 8.14 (32.4) | - | - | 1.93 (3.94) | 0 | 3.19 (2.37) | 0.474 (4.14) |
| Building Footprint (Density) | 10.6 (5.19) | - | - | 17.6 (29.9) | 0 | 43.9 (35.7) | 3.77 (2.39) |
| Mean Building Height | 6.09 (41.6) | - | - | 6.47 (3.19) | 0 | 0 | 0.172 (0.710) |
| Blue Light Pollution | 72.8 (15.1) | - | - | 34.8 (18.0) | 0 | 0 | 5.40 (11.7) |
| Red Light Pollution | 0.0118 (0) | - | - | 0.000189 (0.00465) | 0.718 (8.33) | 0 | 4.78 (3.98) |
| NDVI | 0.203 (0) | - | - | 30.3 (25.8) | 2.81 (6.75) | 0.483 (3.47) | 25.6 (35.7) |
| Forest Proximity | 0.225 (3.91) | - | - | 7.64 (17.6) | 96.5 (84.9) | 51.5 (58.4) | 59.5 (38.1) |
| Model Fit Metrics |  |  |  |  |  |  |  |
| Feature Classes | Q | LQP | Q | LQP | Q | LQP | LQP |
| Regularization Multiplier | 1.5 | 3 | 2.5 | 4.5 | 4.5 | 3 | 5 |
| Null AICc | 699.1862 | 330.3145 | 227.9642 | 2277.736 | 719.6842 | 309.8347 | 2113.725 |
| Model AICc | 688.1845 | 330.3145 | 227.8744 | 2242.165 | 709.5420 | 302.7200 | 2080.960 |
| Number of Observations | 35 | 16 | 11 | 115 | 35 | 15 | 105 |

Appendix S4: Model loadings of all cross-validation runs for each taxon/migratory phenology and spatial scale.

| Taxon | Cross-validation Run | Spatial Scale | Building Height | Forest Proximity | Building Area | Red Light Pollution | Building Perimeter | NDVI | Building Footprint | Blue Light Pollution |
| --- | --- | --- | --- | --- | --- | --- | --- | --- | --- | --- |
| *Chalcophaps indica* | 1 | Fine-scale | 50 | 35.5 | 14.5 | 0 | 0 | 0 | 0 | 0 |
| *Chalcophaps indica* | 2 | Fine-scale | 48.5 | 48.5 | 2.9 | 0 | 0 | 0 | 0 | 0 |
| *Chalcophaps indica* | 3 | Fine-scale | 48.7 | 48.7 | 2.6 | 0 | 0 | 0 | 0 | 0 |
| *Chalcophaps indica* | 4 | Fine-scale | 0 | 91.6 | 5.5 | 0 | 0 | 2.8 | 0 | 0 |
| *Chalcophaps indica* | 5 | Fine-scale | 50 | 50 | 0 | 0 | 0 | 0 | 0 | 0 |
| *Chalcophaps indica* | 1 | Landscape-scale | 0 | 48.9 | 0 | 0 | 0 | 2.2 | 48.9 | 0 |
| *Chalcophaps indica* | 2 | Landscape-scale | 0 | 50 | 0 | 0 | 0 | 0 | 50 | 0 |
| *Chalcophaps indica* | 3 | Landscape-scale | 0 | 36.1 | 0 | 0 | 7.4 | 0 | 56.5 | 0 |
| *Chalcophaps indica* | 4 | Landscape-scale | 0 | 45.6 | 4.4 | 0 | 0 | 0 | 50 | 0 |
| *Chalcophaps indica* | 5 | Landscape-scale | 0 | 84.6 | 0.2 | 0 | 15.2 | 0 | 0 | 0 |
| *Ficedula sp.* | 1 | Fine-scale | 50 | 0 | 0 | 0 | 50 | 0 | 0 | 0 |
| *Ficedula sp.* | 2 | Fine-scale | 50 | 50 | 0 | 0 | 0 | 0 | 0 | 0 |
| *Ficedula sp.* | 3 | Fine-scale | 50 | 50 | 0 | 0 | 0 | 0 | 0 | 0 |
| *Ficedula sp.* | 4 | Fine-scale | 50 | 50 | 0 | 0 | 0 | 0 | 0 | 0 |
| *Ficedula sp.* | 5 | Fine-scale | 49.7 | 49.7 | 0.7 | 0 | 0 | 0 | 0 | 0 |
| *Ixobrychus sp.* | 1 | Fine-scale | 0 | 0 | 0 | 0 | 0 | 0 | 100 | 0 |
| *Ixobrychus sp.* | 2 | Fine-scale | 0 | 0 | 0 | 0 | 0 | 0 | 0 | 0 |
| *Ixobrychus sp.* | 3 | Fine-scale | 0 | 0 | 0 | 0 | 0 | 0 | 100 | 0 |
| *Ixobrychus sp.* | 4 | Fine-scale | 0 | 0 | 68.7 | 0 | 10.9 | 0 | 20.4 | 0 |
| *Ixobrychus sp.* | 5 | Fine-scale | 0 | 0 | 0 | 0 | 0 | 0 | 100 | 0 |
| *Pitta sp.* | 1 | Fine-scale | 2.9 | 0 | 0 | 0 | 0.1 | 0 | 18.5 | 78.6 |
| *Pitta sp.* | 2 | Fine-scale | 0 | 0 | 0 | 0 | 0 | 0 | 25.1 | 74.9 |
| *Pitta sp.* | 3 | Fine-scale | 0 | 0 | 0 | 0 | 4.3 | 0 | 0 | 95.7 |
| *Pitta sp.* | 4 | Fine-scale | 10.2 | 0 | 0 | 0 | 1.1 | 0 | 17.6 | 71.1 |
| *Pitta sp.* | 5 | Fine-scale | 0 | 0 | 0 | 0 | 7.7 | 0 | 0 | 92.3 |
| *Pitta sp.* | 1 | Landscape-scale | 9 | 0 | 1.5 | 0 | 5.8 | 1 | 9.6 | 73.2 |
| *Pitta sp.* | 2 | Landscape-scale | 0.8 | 1.2 | 2.2 | 0.1 | 0 | 0 | 6.1 | 89.7 |
| *Pitta sp.* | 3 | Landscape-scale | 3.1 | 0 | 1.6 | 0 | 11.8 | 0 | 14.7 | 68.8 |
| *Pitta sp.* | 4 | Landscape-scale | 9.7 | 0 | 1.7 | 0 | 11.4 | 0 | 5.9 | 71.3 |
| *Pitta sp.* | 5 | Landscape-scale | 6.8 | 0 | 2.7 | 0 | 11.4 | 0 | 16.9 | 62.2 |
| *Treron sp.* | 1 | Fine-scale | 0 | 99.9 | 0 | 0.1 | 0 | 0 | 0 | 0 |
| *Treron sp.* | 2 | Fine-scale | 0 | 99.1 | 0 | 0.9 | 0 | 0 | 0 | 0 |
| *Treron sp.* | 3 | Fine-scale | 0 | 100 | 0 | 0 | 0 | 0 | 0 | 0 |
| *Treron sp.* | 4 | Fine-scale | 0 | 99.3 | 0 | 0.7 | 0 | 0 | 0 | 0 |
| *Treron sp.* | 5 | Fine-scale | 0 | 100 | 0 | 0 | 0 | 0 | 0 | 0 |
| *Treron sp.* | 1 | Landscape-scale | 0 | 98.1 | 0 | 1.9 | 0 | 0 | 0 | 0 |
| *Treron sp.* | 2 | Landscape-scale | 0 | 87.8 | 0 | 0 | 0 | 12.2 | 0 | 0 |
| *Treron sp.* | 3 | Landscape-scale | 0 | 99.9 | 0 | 0 | 0 | 0.1 | 0 | 0 |
| *Treron sp.* | 4 | Landscape-scale | 0 | 97.7 | 0 | 1.5 | 0 | 0.9 | 0 | 0 |
| *Treron sp.* | 5 | Landscape-scale | 0 | 100 | 0 | 0 | 0 | 0 | 0 | 0 |
| All Migrants | 1 | Fine-scale | 8.9 | 6 | 0.3 | 1.3 | 3.1 | 15.7 | 40.9 | 24 |
| All Migrants | 2 | Fine-scale | 7.4 | 13.9 | 0 | 0.4 | 3.5 | 8.3 | 41.9 | 24.5 |
| All Migrants | 3 | Fine-scale | 9.2 | 11.7 | 0.2 | 3.5 | 8.3 | 17.6 | 26.7 | 22.6 |
| All Migrants | 4 | Fine-scale | 6.2 | 9 | 0 | 2.4 | 2.3 | 12.4 | 39.4 | 28.4 |
| All Migrants | 5 | Fine-scale | 0 | 8.8 | 0.7 | 0 | 7.3 | 29.3 | 43.1 | 10.9 |
| All Migrants | 1 | Landscape-scale | 12.5 | 19.1 | 1 | 0 | 0.9 | 22.9 | 12 | 31.5 |
| All Migrants | 2 | Landscape-scale | 8.5 | 3.8 | 2.6 | 0 | 5.5 | 23.1 | 16.1 | 40.4 |
| All Migrants | 3 | Landscape-scale | 3.2 | 3.4 | 0.9 | 0 | 0.9 | 44.2 | 21.6 | 25.9 |
| All Migrants | 4 | Landscape-scale | 3.6 | 6.4 | 0.6 | 0 | 0.3 | 33.4 | 25.2 | 30.5 |
| All Migrants | 5 | Landscape-scale | 4.9 | 6.5 | 0.9 | 0 | 1.8 | 28 | 12.3 | 45.6 |
| All Residents | 1 | Fine-scale | 2 | 82.3 | 0 | 1 | 1 | 2 | 5.9 | 5.8 |
| All Residents | 2 | Fine-scale | 1.2 | 72.1 | 0.6 | 0.8 | 0.9 | 9.5 | 7.1 | 7.7 |
| All Residents | 3 | Fine-scale | 1.5 | 87 | 0.4 | 3.6 | 5.3 | 2 | 0 | 0.3 |
| All Residents | 4 | Fine-scale | 1.9 | 74.3 | 1 | 0.2 | 6.9 | 8.7 | 0.1 | 6.9 |
| All Residents | 5 | Fine-scale | 1.7 | 90.1 | 0 | 0.1 | 0.2 | 2.5 | 2.4 | 2.9 |
| All Residents | 1 | Landscape-scale | 0 | 69.4 | 0 | 5.9 | 0 | 24.3 | 0.2 | 0.2 |
| All Residents | 2 | Landscape-scale | 0 | 64 | 0.6 | 2.6 | 0.6 | 28.3 | 2.2 | 1.8 |
| All Residents | 3 | Landscape-scale | 0 | 65.9 | 0.8 | 0 | 0.9 | 18.9 | 1.8 | 11.6 |
| All Residents | 4 | Landscape-scale | 0.7 | 37.3 | 0 | 9.3 | 0.7 | 28.3 | 14.1 | 9.5 |
| All Residents | 5 | Landscape-scale | 0.1 | 61.5 | 0.2 | 5.7 | 0.2 | 27.9 | 0.2 | 4.2 |

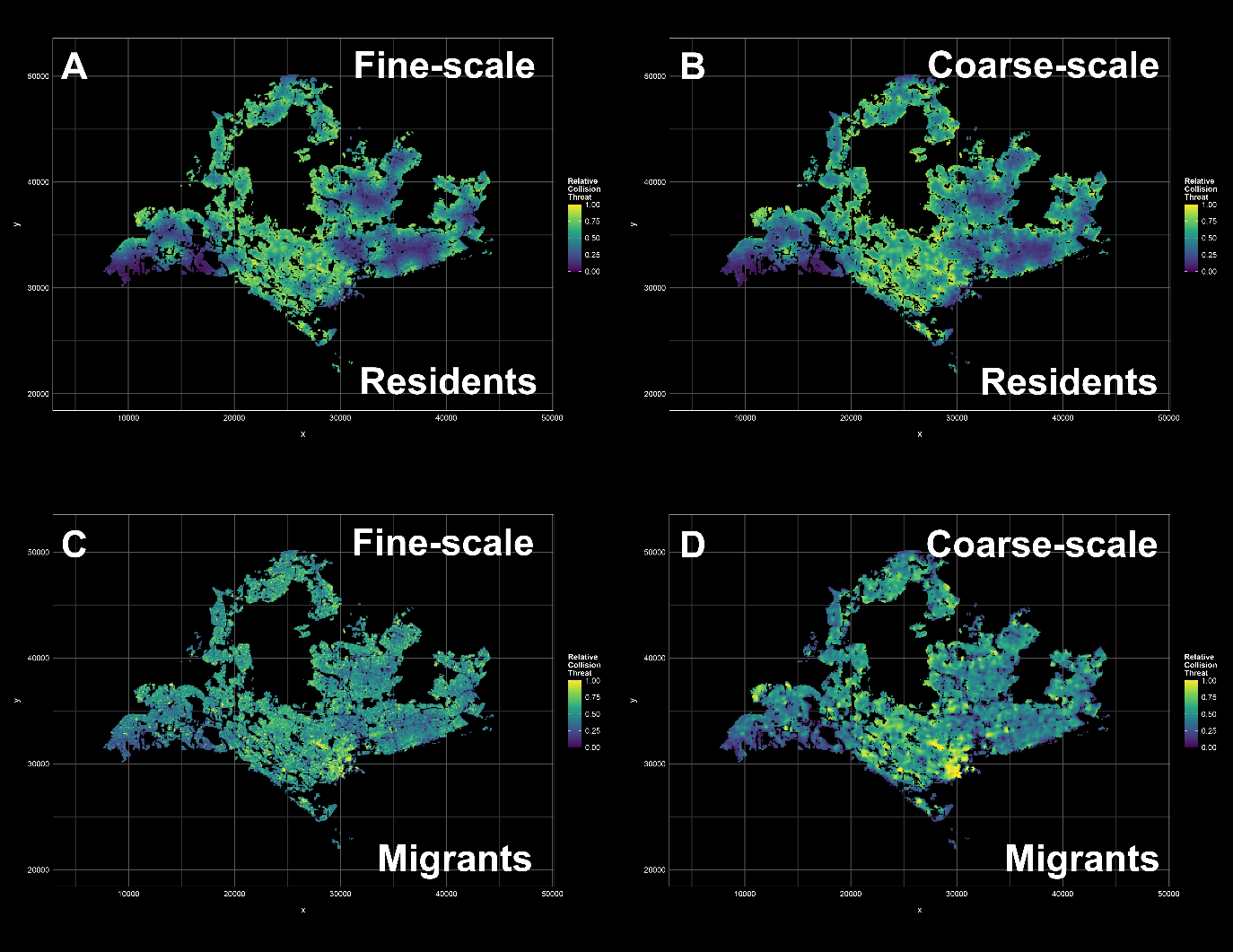

Appendix S5: Building collision risk maps for (A and B) all resident species, and (C and D) all migratory species, for both fine-scale (A and C) and coarse-scale (B and D) models.

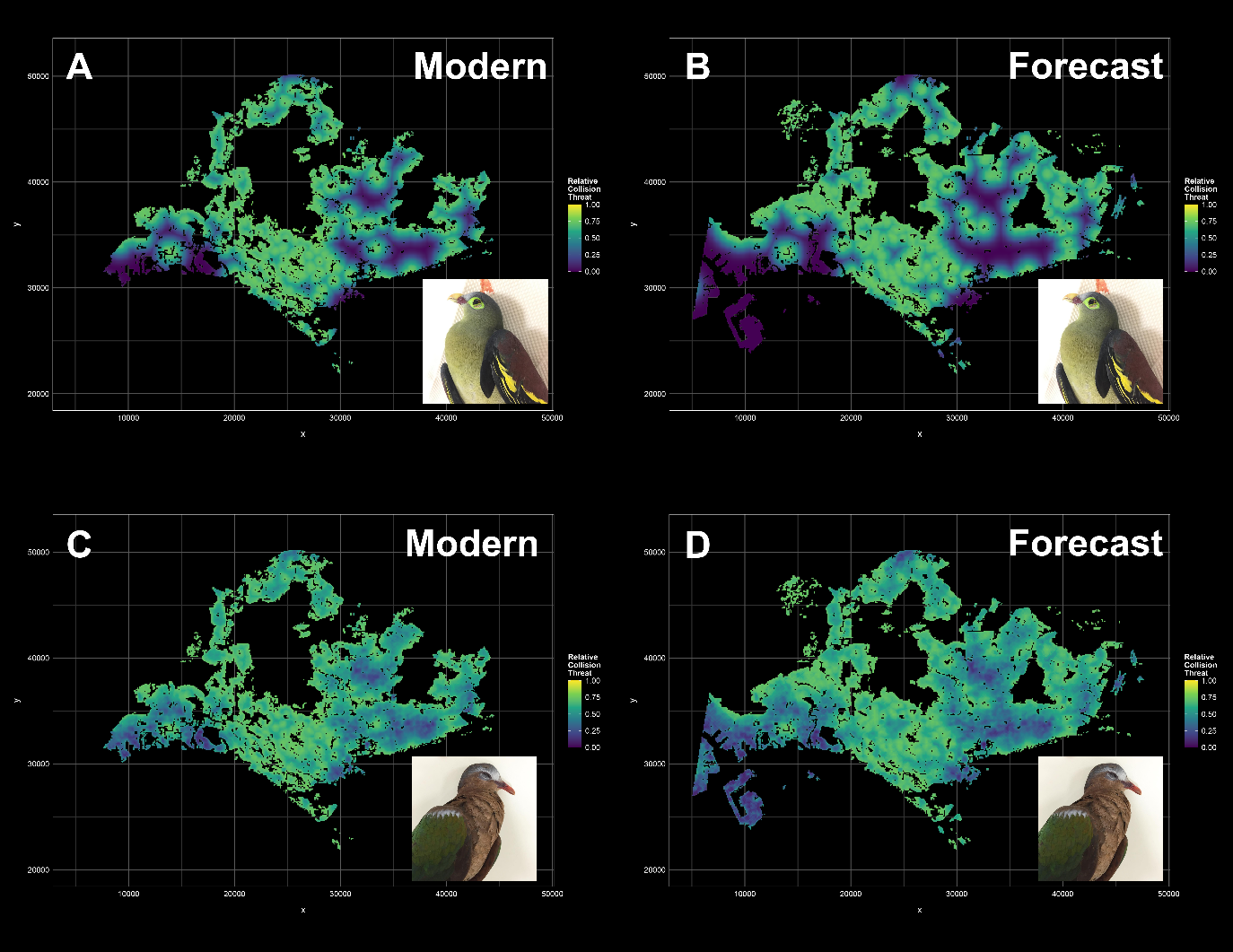

Figure S6: Modern-day and forecast building collision risk maps for Green Pigeons (*Treron sp.*; panels A and B) and Asian Emerald Doves (*Chalcophaps indica*; panels C and D), based on coarse-scale predictor variables, showing a similar collision risk distribution to the fine-scale models.

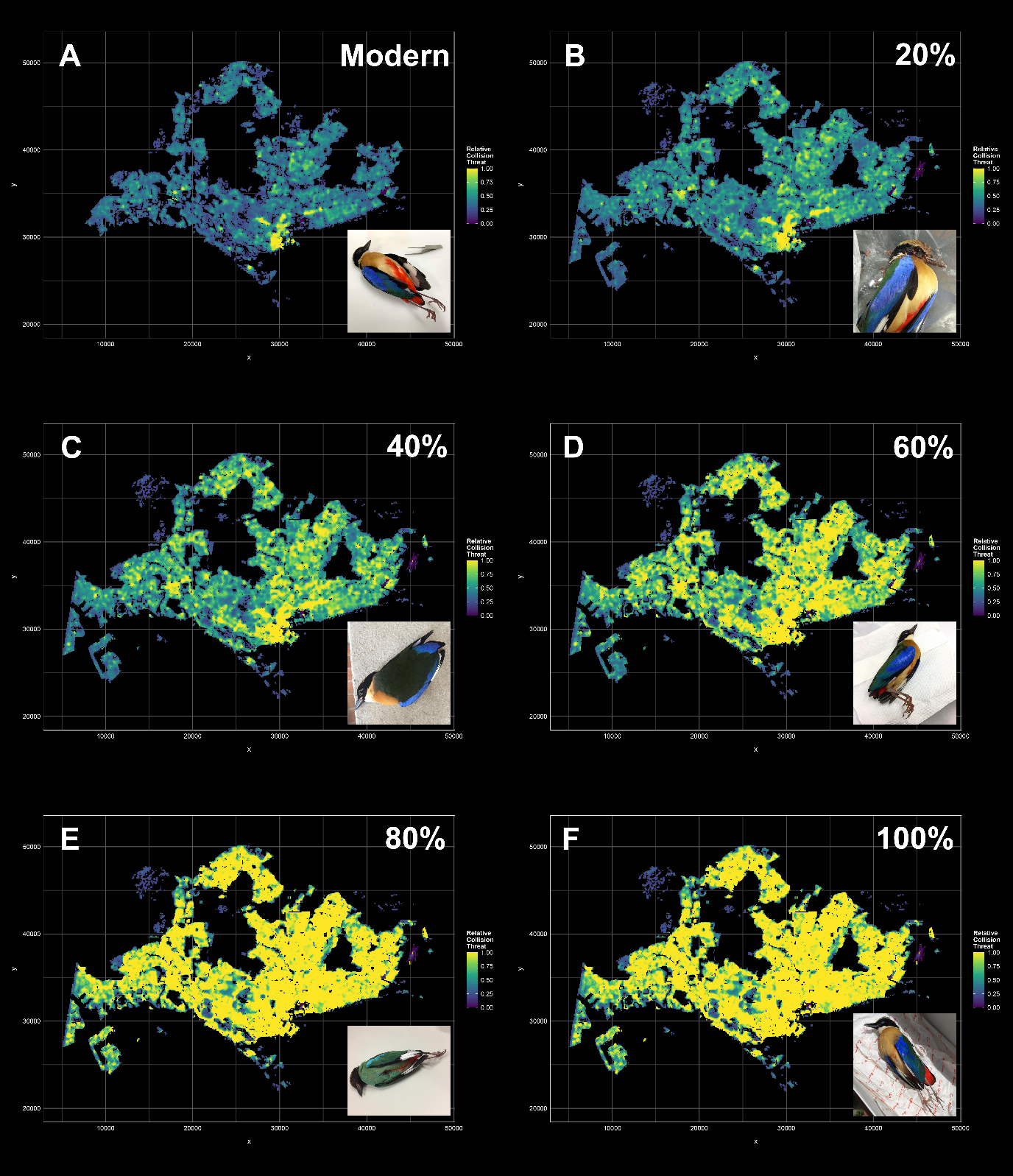

Figure S7: Coarse-scale collision risk maps for the True Pittas (*Pitta sp.*), for increasing amounts of projected future blue light pollution. These scenarios correspond with (A) 0% (modern-day), (B) 20%, (C) 40%, (D) 60%, (E) 80%, and (F) 100% increases in blue light output relative to present-day levels of red streetlight pollution.

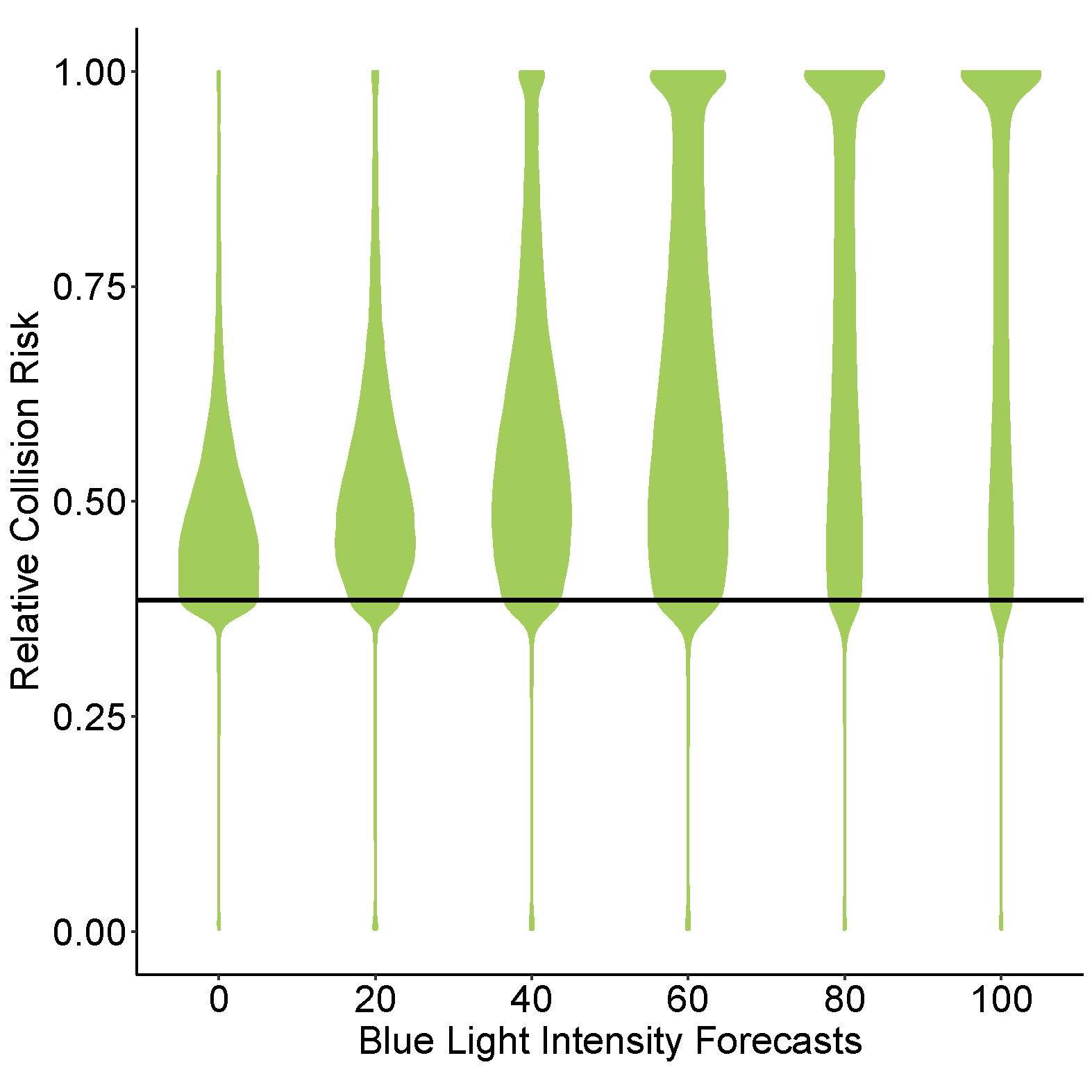

Figure S8: Violin plots showing how projected future increases in nocturnal blue light pollution are likely expected to affect the landscape-wide distribution of relative building collision risks for the True Pittas. The results indicate that an increase in blue light pollution exceeding 40% of present-day red streetlight pollution is likely to result in a notable upward shift in the distribution of relative collision risk.

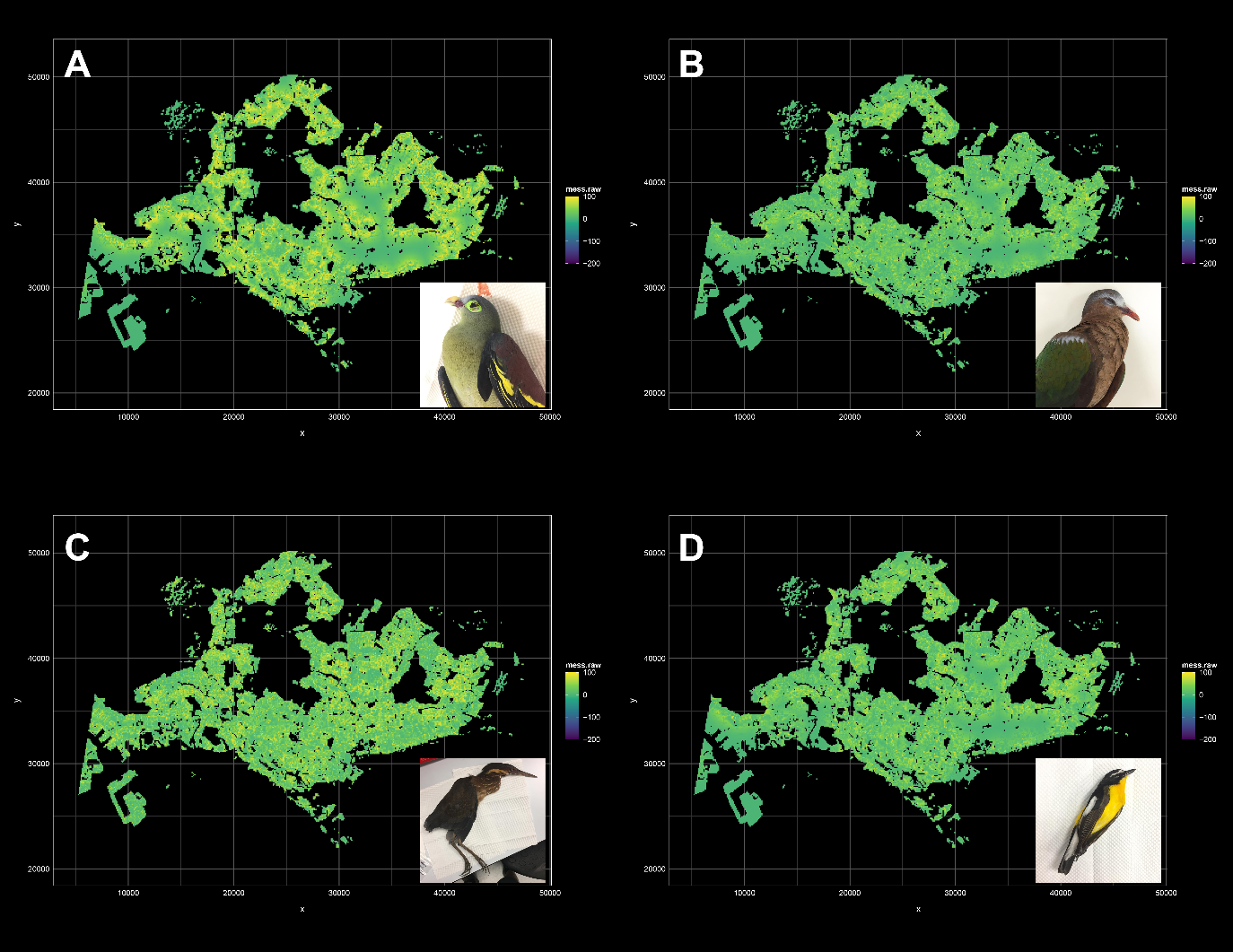

Figure S9: Fine-scale multivariate environmental similarity surface (MESS) maps for (A) Green Pigeon, (B) Asian Emerald Dove, (C) Bittern, and (D) *Ficedula* flycatcher collision risk projections, showing that the prediction dataset largely falls within the parameter space of the training dataset.

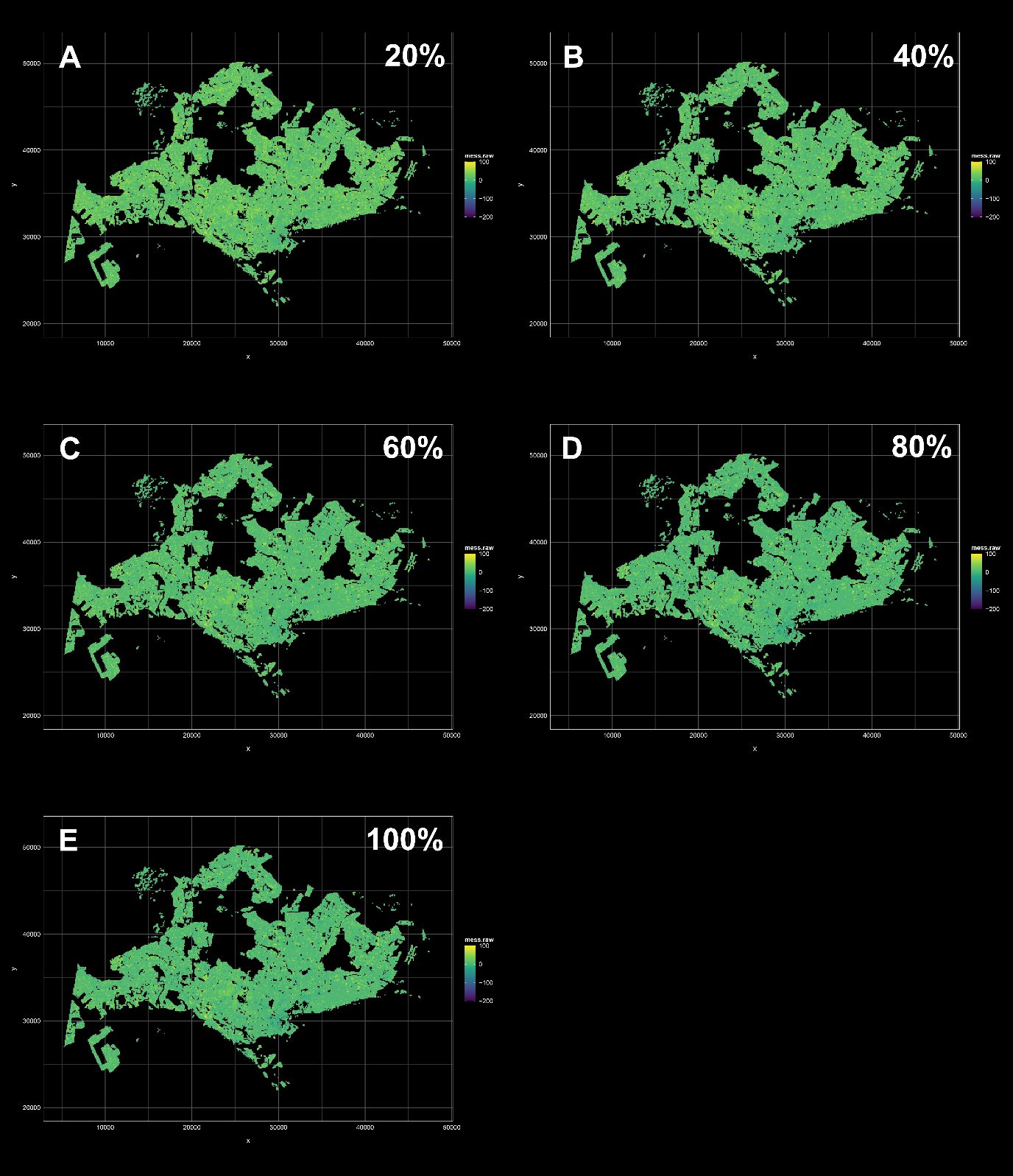

Figure S10: Fine-scale multivariate environmental similarity surface (MESS) maps for the True Pittas (*Pitta sp.*), for five projected scenarios of future nocturnal blue light pollution, corresponding with (A) 20%, (B) 40%, (C) 60%, (D) 80%, and (E) 100% increase in blue light pollution relative to present-day red streetlight pollution.

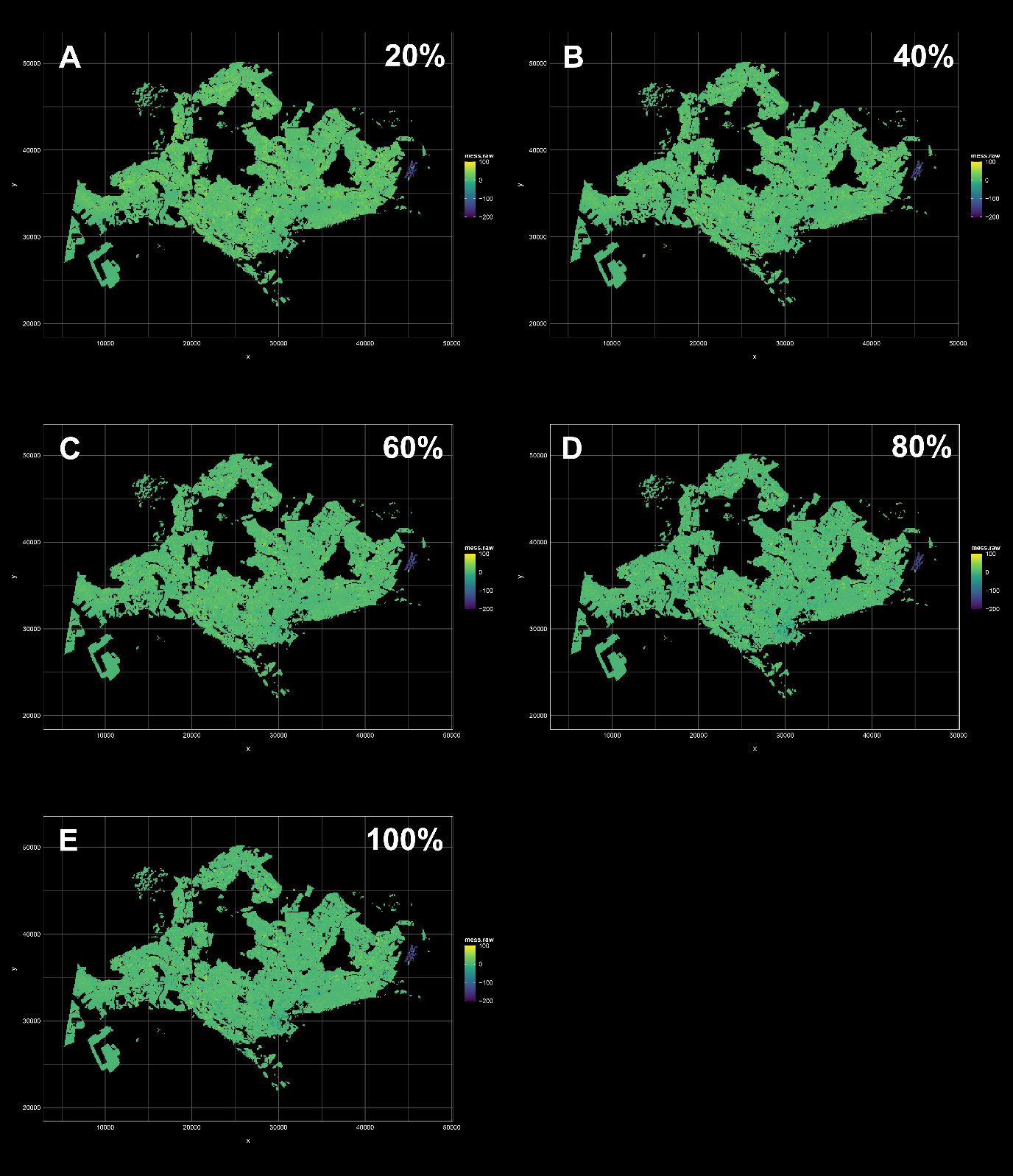

Figure S11: Fine-scale multivariate environmental similarity surface (MESS) maps for all resident species, for five projected scenarios of future nocturnal blue light pollution, corresponding with (A) 20%, (B) 40%, (C) 60%, (D) 80%, and (E) 100% increase in blue light pollution relative to present-day red streetlight pollution.

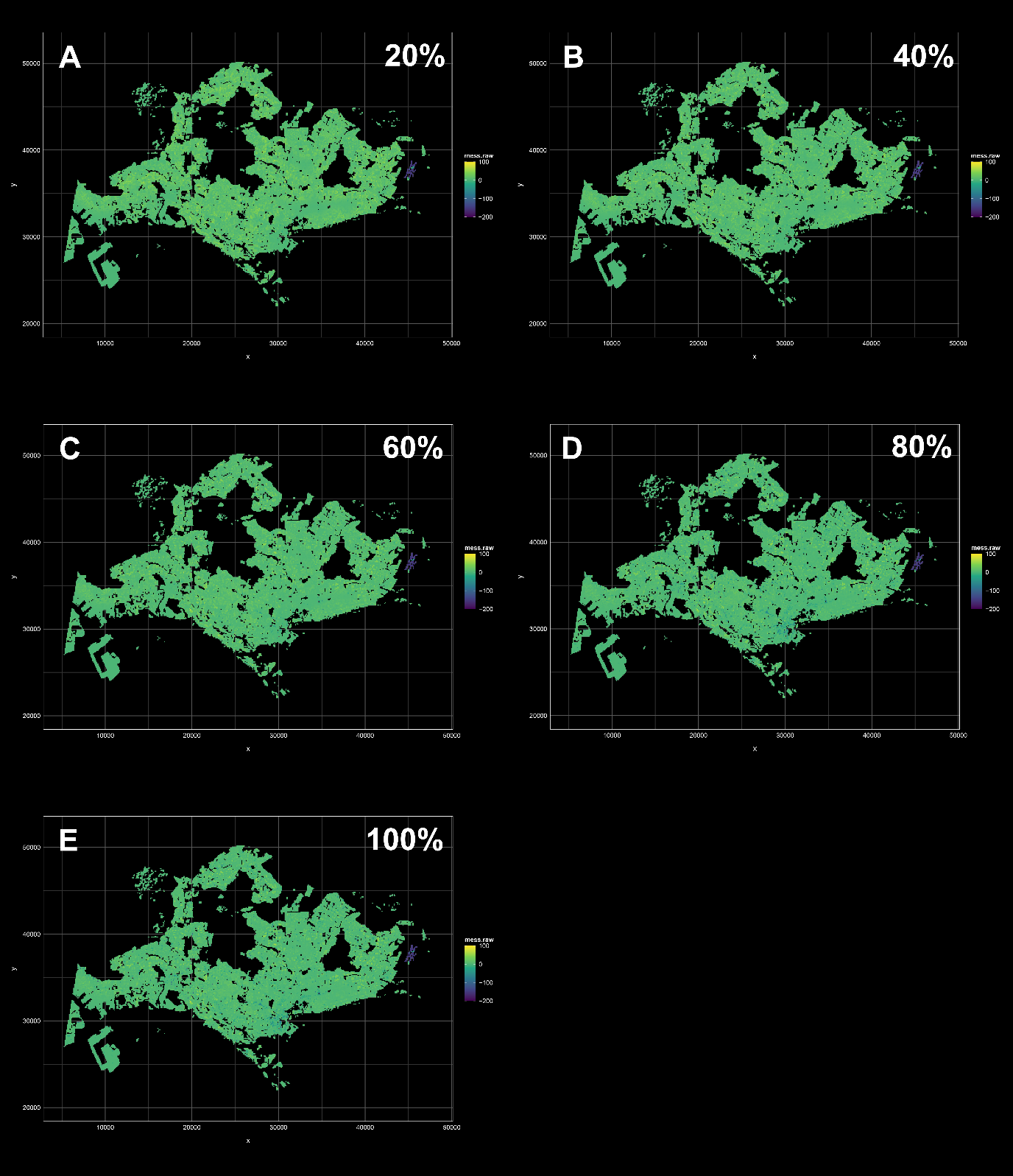

Figure S12: Fine-scale multivariate environmental similarity surface (MESS) maps for all migratory species, for five projected scenarios of future nocturnal blue light pollution, corresponding with (A) 20%, (B) 40%, (C) 60%, (D) 80%, and (E) 100% increase in blue light pollution relative to present-day red streetlight pollution.

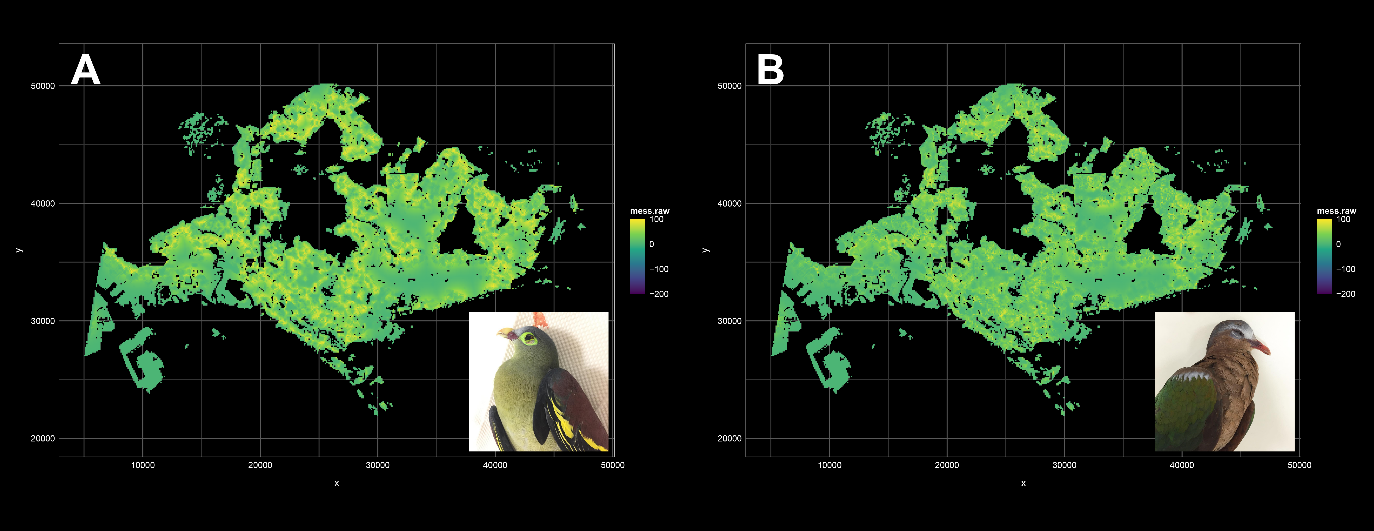

Figure S13: Coarse-scale multivariate environmental similarity surface (MESS) maps for (A) Green Pigeons, (B) Asian Emerald Doves.

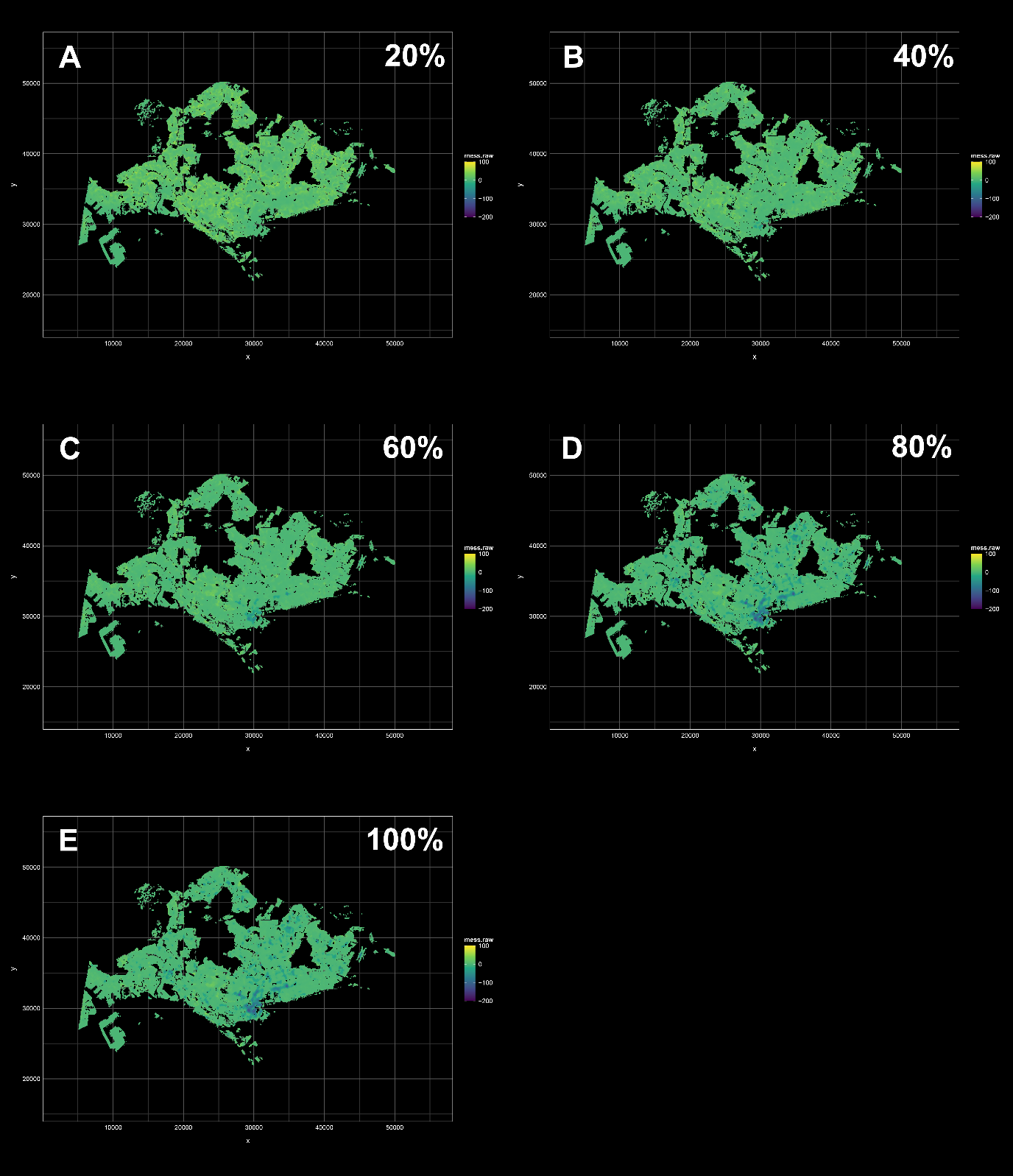

Figure S14: Coarse-scale multivariate environmental similarity surface (MESS) maps for the True Pittas (*Pitta sp.*), for five projected scenarios of future nocturnal blue light pollution, corresponding with (A) 20%, (B) 40%, (C) 60%, (D) 80%, and (E) 100% increase in blue light pollution relative to present-day red streetlight pollution. The maps show that at models with high projected levels of blue light pollution are likely to exhibit lower suitabilities, indicating a greater degree of extrapolation from the training dataset.

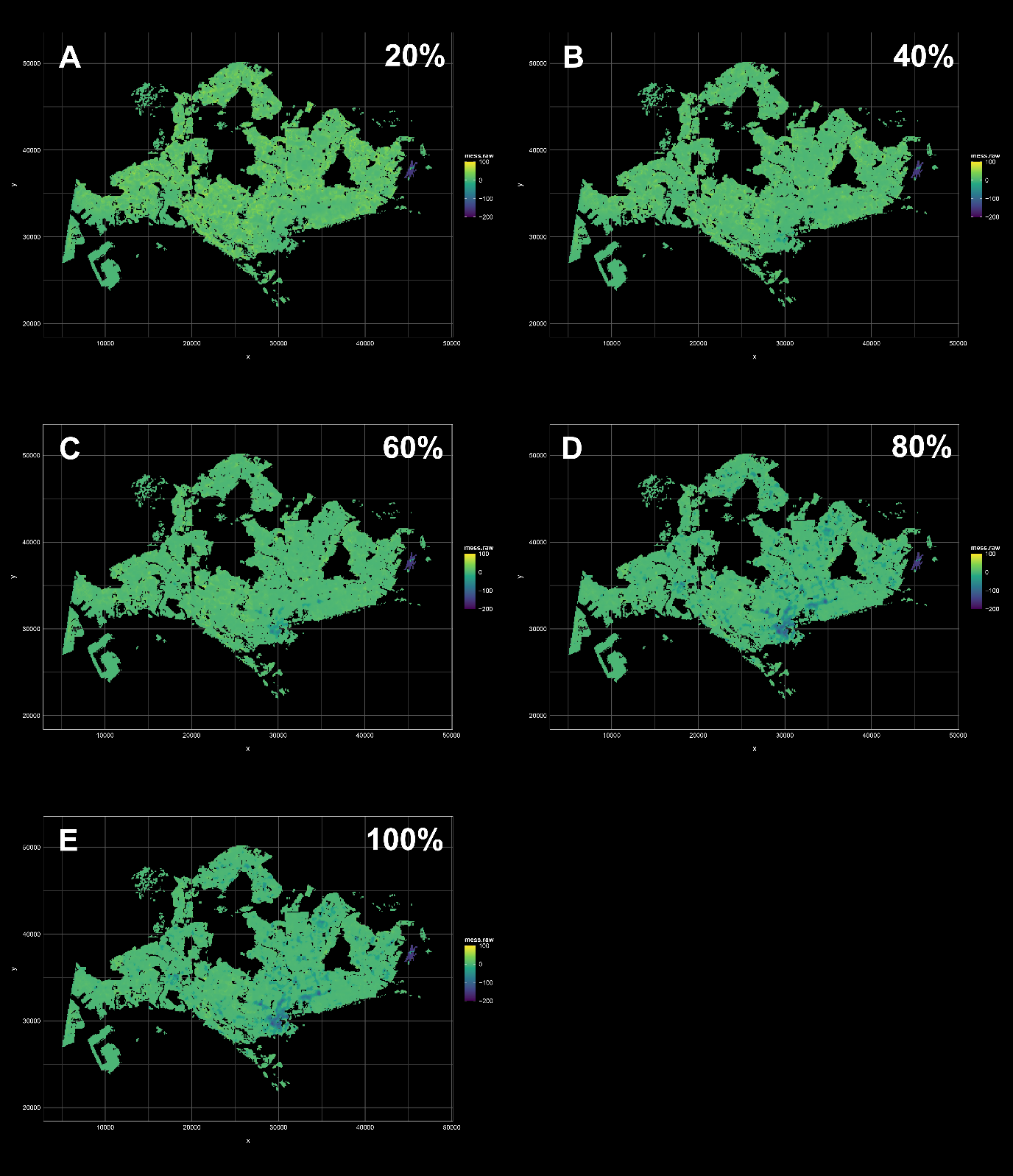

Figure S15: Coarse-scale multivariate environmental similarity surface (MESS) maps for all resident species, for five projected scenarios of future nocturnal blue light pollution, corresponding with (A) 20%, (B) 40%, (C) 60%, (D) 80%, and (E) 100% increase in blue light pollution relative to present-day red streetlight pollution. The maps show that at models with high projected levels of blue light pollution are likely to exhibit lower suitabilities, indicating a greater degree of extrapolation from the training dataset. The anomalously large Changi Airport building in the east of Singapore is also likely to fall outside the parameter range of the training dataset.

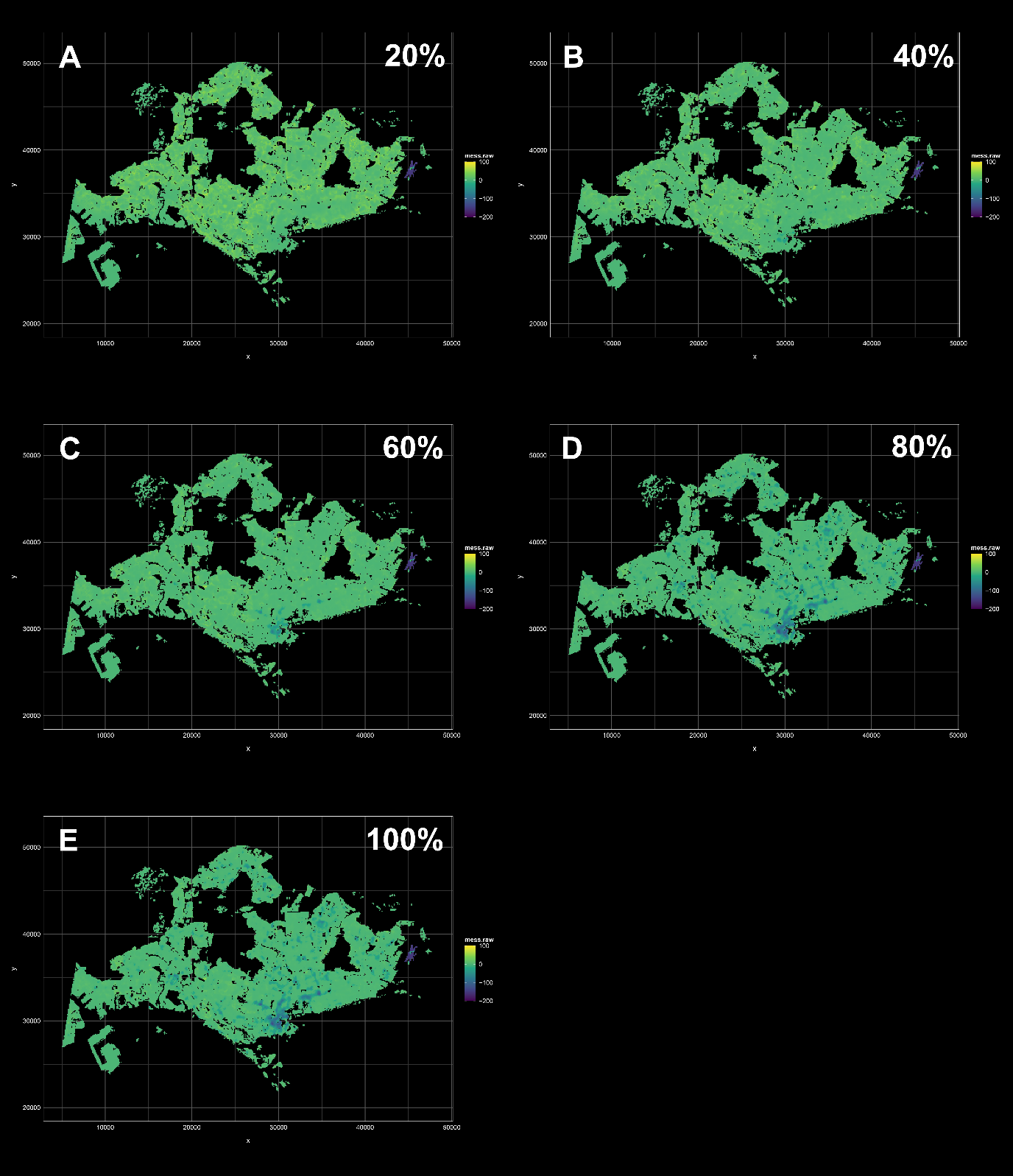

Figure S16: Coarse-scale multivariate environmental similarity surface (MESS) maps for all migrant species, for five projected scenarios of future nocturnal blue light pollution, corresponding with (A) 20%, (B) 40%, (C) 60%, (D) 80%, and (E) 100% increase in blue light pollution relative to present-day red streetlight pollution. The maps show that at models with high projected levels of blue light pollution are likely to exhibit lower suitabilities, indicating a greater degree of extrapolation from the training dataset. The anomalously large Changi Airport building in the east of Singapore is also likely to fall outside the parameter range of the training dataset.
